# Clustered Regularly Interspaced Short Palindromic Repeats-Cas system regulates *Salmonella* virulence

**DOI:** 10.1101/2023.12.12.571267

**Authors:** Nandita Sharma, Ankita Das, Abhilash Vijay Nair, Palash Sethi, Vidya Devi Negi, Dipshikha Chakravortty, Sandhya Amol Marathe

**Author notes:** These authors contributed equally to this study (Nandita Sharma, Ankita Das). Correspondence to: S. A. Marathe.

## Abstract

**Objectives:** Investigating the type 1-E CRISPR-Cas-mediated regulation of *Salmonella* pathogenesis.

**Methods:** We assessed the pathogenicity of the wild-type and CRISPR-Cas knockout strains using infection models. The mechanisms were elucidated using antimicrobial assays and expression analysis.

**Results:** CRISPR-Cas knockout strains were defective in invasion and proliferation in intestinal epithelial cells and macrophages. However, proliferation defects were not observed in the Gp91^phox-/-^ macrophages, suggesting the system’s role in antioxidant defence. The knockout strains show hampered colonization in *in-vivo* infection models, possibly due to increased sensitivity against innate immune barriers like antimicrobial peptides, complement proteins and oxidative stress. The expression studies of various virulence regulators: *pmr* genes, anti-oxidant genes, SPI-1 and SPI-2 encoded master regulators, and effectors showed repressed expression in the knockout strains. Some of these genes could be directly regulated by the CRISPR-spacers owing to partial complementarity between the sequences.

**Conclusion:** Overall, our study shows that the CRISPR-Cas system positively regulates *Salmonella* pathogenesis by regulating the expression of different virulence factors.

## Introduction

The bacterial adaptive immune system, Clustered Regularly Interspaced Short Palindromic Repeats (CRISPR), and CRISPR-associated (Cas) endonucleases, acts against invading mobile genetic elements. Besides their canonical functions, the CRISPR-Cas systems regulate the physiology and virulence of various bacteria, including *Streptococcus*, *Enterobacter*, and *Salmonella* [1,2]. For example, Cas 9 of *Francisella* targets bacterial lipoprotein (BLP) mRNA, thereby subverting the host immune response [3]. Cas9 aids the invasion and intracellular survival of *Campylobacter* and *Neisseria* [4,5]. In *Legionella,* Cas2 is required to infect amoebae[6]. The CRISPR-RNA can also work independently of Cas proteins and regulate bacterial virulence. In *Listeria monocytogenes,* crRNA governs its virulence in the absence of *cas* genes [7]. Similarly, in *Streptococcus agalactiae,* CRISPR regulates virulence independently of Cas [8]. The type I-F CRISPR-Cas system of *Pseudomonas aeruginosa* regulates biofilm and virulence by targeting *lasR* transcript [9]. Further, the computational analysis predicted the role of the type 1-E system of *E. coli* in endogenous gene regulation [10]. However, the mechanistic details are yet to be fully investigated.

Many proto-spacers of the CRISPR-Cas system in *Salmonella* were traced on the chromosome instead of its general target phages and plasmids but the target genes were unidentified [11]. A few studies have suggested the CRISPR-Cas system’s role in modulating *Salmonella enterica* subspecies *enterica* serovar Typhi’s (*S*. Typhi) intracellular survival and biofilm formation[12]. The expression of *cas7* was detected in human macrophages infected by *S*. Typhi. The transcriptome profile of bacteria displayed altered expression of *cas* genes in clinical *S.* Typhi samples, suggesting the role of the type I-E CRISPR-Cas system during *Salmonella* infection [13]. A recent study by Cui *et al.* on *Salmonella enterica* subspecies *enterica* Enteritidis (*S*. Enteritidis) correlates the importance of the CRISPR-Cas system in regulating quorum sensing, biofilm formation, and bacterial invasion into the host [14]. The study indicates the role of Cas3 nuclease in *S*. Enteritidis in regulating virulence and biofilm formation. The results of the study hint at the regulation of invasion and QS genes by Cas3. Another relevant study was performed by Stringer *et al.* ChIP seq analysis confirmed 236 crRNA and Cascade-binding sites in *S. enterica* subsp. *enterica* serovar Typhimurium (*S*. Typhimurium) [15]. These sites coincide with virulence gene loci - *sseA, bcsA, iroC, entE, entF, sptP,* possibly suggesting the regulation of pathogenic traits by the CRISPR-Cas system. Furthermore, the CRISPR array and Cas genes exert regulatory control over the expression of a diverse array of genes associated with biofilm formation in *Salmonella*, like *csgA*, *csgD*, *yddX*, *bcsA*, and *bcsC*. This regulation distinctly governs the processes of surface and pellicle biofilm formation [16].

The infection biology of *Salmonella* is sequentially orchestrated into five steps – attachment to the intestinal epithelium, invasion in intestinal epithelial cells, proliferation inside macrophages, systemic dissemination in various organs, persistence in the gallbladder, and transmission by fecal shedding. The outcome of *Salmonella* infection is determined by the virulence of the infecting *Salmonella*, the ability of the host to mount an adequate immune response, and the ability of the host to destroy the pathogen. Typically, the infection process begins *via* oral intake of *Salmonella-*contaminated food. The intestinal adhesion and invasion mark the beginning of *Salmonella* pathogenesis [17]. The virulence of *Salmonella* is influenced by the pathogenicity determinants encoded on the pathogenicity islands, SPI-1 (regulates invasion phase) and SPI-2 (regulates proliferation phase)[18]. The type III secretion apparatus aids in transferring effector proteins into host cells. Various environmental clues in the intestine induce the expression of the *Salmonella* pathogenicity island 1 (SPI-1) transcriptional regulator, HilD. HilD binds upstream of the master regulator, *hilA*, thereby counteracting the H-NS-mediated repression of the *hilA* promoter [19]. Following the activation of *hilA*, the structural genes and effector proteins encoded by SPI-1 get activated[19]. Following the internalization of bacteria, the *Salmonella*-containing vacuole (SCV), induces the expression of the SPI-2-encoded SsrAB system [19]. The SsrA kinase phosphorylates the key regulatory factor of SPI-2, SsrB, which in turn activates the expression of SPI-2 encoded effector proteins[20]. Multiple other SPI-2 effectors govern the replication and survival of the bacteria within the host.

Building upon these findings, we decided to investigate the involvement of the type IE CRISPR-Cas system in regulating *Salmonella’s* virulence at different infection stages. The CRISPR-Cas knockout strains were attenuated in virulence. Thus, we checked for their ability to evade the host defenses through the regulation of important virulence genes.

## Material and Methods

### Bacterial strains, nematode, and culture conditions

The wildtype *Salmonella enterica* serovar Typhimurium strain 14028s was used as the parent strain in this study. The wildtype, knockout (Δ*crisprI,*Δ*crisprII,* ΔΔ*crisprI crisprII* and Δ*cas op)* and complement (Δ*crisprI+pcrisprI* and Δ*crisprII+pcrisprII*) strains were cultured in Luria-Bertani ( LB, Himedia) with appropriate antibiotics. The mCherry fluorescent derivatives of wildtype, CRISPR-Cas knockout strains, and *Escherichia coli* OP50 were obtained by electro-transformation of a pFPV-mCherry plasmid, a kind gift from Prof. Dipshikha Chakravorty. Bacterial cultures were routinely maintained on LB-Agar media along with their corresponding antibiotics (Supplementary Table 1). A wildtype N2 strain of *Caenorhabditis elegans* was routinely maintained at 25°C on a nematode growth medium (NGM) agar plate with *E. coli* OP50 as a food source.

### Eukaryotic cell lines and growth conditions

The murine macrophage-like cell line RAW 264.7, and a colorectal adenocarcinoma cell line with epithelial morphology (HT-29) cells were obtained from NCCS, Pune. These cells were grown in Dulbecco′s modified minimum essential medium (Gibco) and Roswell Park Memorial Institute 1640 media (Sigma Aldrich), respectively, with 10% fetal bovine serum (FBS, Himedia) at 37°C temperature in the presence of 5% CO_2_. RAW 264.7 cells were activated with 10 ng/mL Lipopolysaccharide (LPS) from *E.coli* (Sigma) for 24 h. For polarizing the HT-29 cells, RPMI was supplemented with 2 mM glutaMAX^TM^ for 15 days (Gibco).

Murine peritoneal macrophages were harvested as described previously [21]. BALB/c, C57BL/6 and gp91^phox−/−^ mice were injected intraperitoneally with 1 mL of 3% thioglycollate (Himedia). After four days, the mice were sacrificed by cervical dislocation, 5 mL of ice-cold RPMI media was injected into the peritoneal cavity of mice, and peritoneal macrophages were withdrawn using a syringe. Cells were centrifuged and suspended in RPMI media with 10% FBS.

### Percentage phagocytosis/invasion assay

Bacterial phagocytosis and invasion were estimated by a gentamicin protection assay in macrophages (RAW264.7 and peritoneal macrophages) and intestinal epithelial cell lines (HT-29 non-polarized and polarized cell lines), respectively. 1.5 to 3×10^5^ cells were seeded into 24-well plates and incubated at 37°C with 5% CO_2_ for 24 h. RAW 264.7 and peritoneal macrophages were infected with stationary phase cultures of wildtype, Δ*crisprI,* Δ*crisprII,* Δ*cas op,* and ΔΔ*crisprI crisprII* knockout strains and their respective complement strains Δ*crisprI+*p*crisprI* and Δ*crisprII+*p*crisprII* at a multiplicity of infection (MOI) 5. The HT-29 cells were infected with the log phase culture (when the SPI-1 system is active) of the strains at MOI 10. The infected cells were incubated at 37°C with 5% CO_2_ for 30 min. Next, the cells were washed thrice with phosphate-buffered saline (PBS) and subjected to 100 μg/mL of gentamicin treatment for 1 h. The cells were washed again with PBS and lysed with 0.5 mL of 0.1% Triton X-100 (Sigma). The number of viable intracellular bacteria was determined by plating the lysates onto LB agar supplemented with appropriate antibiotics. Percentage phagocytosis/invasion was determined using the following formula:

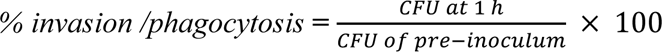

### Intracellular proliferation assay

The protocol for intracellular proliferation is similar to that mentioned above. The cells were infected with the stationary phase (RAW 264.7 cells and peritoneal macrophages) or log phase (HT-29 cells) bacterial cultures at MOI of 5 and 10, respectively. The cells were lysed at 2 h and 16 h post-infection. The lysates were plated onto LB agar supplemented with antibiotics to obtain CFU at 2 h and 16 h. The fold proliferation 16 to 2 h was determined using the following formula:

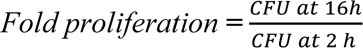

### *In-vivo* infection assay

For infection studies, 6-8 weeks old BALB/c mice weighing 20-22 g raised in Central Animal Facility, Indian Institute of Science (IISc), Bangalore were used as per the guidelines of the Institutional Animal Ethics Committee at the IISc, Bangalore, India. Five mice in five sets were orally gavaged with 10^7^ bacterial cells of wild type, Δc*risprI,* Δ*crisprII,* Δ*cas op,* and ΔΔ*crisprI crisprII* knockout strains. After 3 days post-infection, reticuloendothelial organs like the spleen, liver, Peyer′s patch (PP), and mesenteric lymph nodes (MLN) were aseptically isolated, weighed, and homogenized in 0.5 mL of sterile PBS using a bead-beater (Bio spec products, USA). Serial dilutions of the homogenate were plated onto *Salmonella Shigella* agar (SS agar, Himedia) containing appropriate antibiotics to obtain CFU per gram weight for each organ.

#### Cytokine analysis

The blood from control and infected mice was collected on the 4^th^ day by the retro-orbital bleeding method, and the sera was separated once the blood was clotted. The concentration of IFN-γ, IL-4, and IL-10 in the pooled sera of each set was estimated using a Thermo Fisher Scientific kit as per the manufacturer’s instructions.

### Bacterial colonization assay in *C. elegans*

The mCherry-tagged bacterial strains were grown overnight in LB broth at 37°C and lawns were prepared by spreading 200 µL of overnight bacterial culture on modified NGM agar (Agar-1.7 g, NaCl-0.3 g, peptone-0.05 g). To measure the intestinal colonization of the test strains in *C. elegans*, the synchronized L4 larvae were exposed to fluorescently-tagged (m-Cherry) strains of wildtype, Δ*crisprI,* Δc*risprII,* Δ*cas op,* ΔΔ*crisprI crisprII*, and *E. coli* OP50. After 24 h, the worms were anesthetized with 25 mM levamisole (Sigma), washed thrice with M9 buffer (KH_2_PO_4_, Na_2_HPO_4_, 5 g NaCl and 1 M MgSO_4_), and treated with 80 μg/mL of gentamicin for 1 h followed by treatment with 25 μg/mL of gentamicin for 30 min. Finally, the worms were washed with M9 buffer and lysed with 0.2% Triton X-100 (Sigma) in a tissue lyser LT (Qiagen, India). The lysates were serially diluted and plated on LB-agar containing ampicillin to estimate bacterial burden.

### Antimicrobial peptide killing assay

Overnight-grown bacterial cultures were subcultured at a ratio of 1:40 in Luria broth and incubated at 37°C until the OD_600nm_ reached 0.3-0.4. 10^5^ bacterial cells were treated with polymyxin B (0.5 μg/mL, Himedia), and protamine sulfate (0.5µg/mL, PROTA), in TN (0.5% tryptone and 0.5% NaCl) media for 1 h at 37°C with slight agitation. Following the incubation, the mixture was plated onto LB-agar plates supplemented with appropriate antibiotics. Percentage survival was calculated with respect to the untreated samples.

### Serum sensitivity

Bacterial strains were grown overnight in Luria broth, and 10^7^ bacterial cells from an overnight culture were incubated in 20% FBS (Himedia) for 2 h at 37°C with slight agitation. Serial dilutions of these cultures were plated onto LB agar supplemented with antibiotics to determine the CFU. The percentage survival was calculated using the following formula:

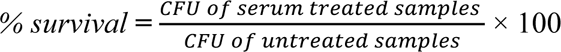

### Measurement of intracellular reactive oxygen species (ROS)

LPS activated RAW 264.7 cells were infected with wildtype, Δ*crisprI,* Δ*crisprII,* Δc*as op, and* ΔΔ*crisprI crisprII* knockout strains at MOI 5 as described in the above sections. The intracellular ROS was determined using an oxidant-sensitive probe 2′,7′-dichlorodihydrofluorescein diacetate (H_2_DCFDA, Sigma) 6h post-infection. The culture supernatant was replaced with DMEM media, supplemented with a 5 µM concentration of H_2_DCFDA, followed by incubation at 37°C, 5% CO_2_ for 30 min. The cells were washed with sterile PBS, and fluorescence intensity was measured at (λ_ex_) of 485 nm and (λ_em_) at 535 nm using Fluoroskan (Thermo Scientific).

### Measurement of extracellular reactive nitrogen species (RNS)

Extracellular nitrite was measured as described previously [22]. RAW 264.7 cells were infected, as described in the above section. 50 µL the extracellular media were collected from cells infected with wildtype, Δ*crisprI,* Δ*crisprII,* Δ*cas op, and* ΔΔ*crisprI crisprII* knockout strains at 16 h post-infection and subjected to nitrite estimation by Griess reagents. To the culture media, 50 µL each of 1% sulphanilamide (made in 5% phosphoric acid), and 0.1% NED (N-1-naphthyl ethylenediamine dihydrochloride) (Himedia) were added and incubated in the dark at room temperature for 15 min. OD_545nm_ was measured within 30 min of the appearance of a purple-coloured product.

### Bacterial sensitivity to hydrogen peroxide

Overnight grown bacterial cultures of wildtype, Δ*crisprI,* Δ*crisprII,* Δ*cas op,* ΔΔ*crisprI crisprII,* Δ*crisprI+*p*crisprI* and Δ*crisprII+*p*crisprII* strains were treated with 1 mM H_2_O_2_ in Muller Hinton (MH, Himedia, pH- 5.4) media for 2 h. The bacterial suspensions were plated onto LB-agar plates containing appropriate antibiotics to determine Colony Forming Units (CFU).

#### Priming assay with hydrogen peroxide

The overnight grown bacterial cultures were subcultured in MH Media and grown to OD_600nm_ ∼ 0.4. For one set, the bacteria were exposed to 0.1 mM H_2_O_2_ (priming) for 30 min in the dark at 37°C with shaking. The H_2_O_2_ was removed by centrifugation at 5000 x g for 10 min, and cells were allowed to recover for 2 h. The other set was left untreated. Equal amounts (10^7^) of bacteria from each set were subcultured in MH media. The bacteria were incubated with 0 mM and 1 mM (trigger), of H_2_O_2_ at 37°C in the dark at 100 rpm. After 8 h, the bacterial growth was determined by measuring the OD_600nm_ using Multiskan GO (Thermo Scientific, USA).

The percentage survival was calculated using the following formula:

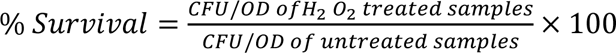

### RNA isolation and quantitative real-time (q-RT) PCR

Bacterial strains grown overnight were subcultured at a ratio of 1:100 in LB (SPI-I inducing condition) and F media (pH= 5.4, SPI-2 inducing condition). The bacterial cells were incubated at 37°C and 150 rpm for 8 h and 4 h, respectively. At the end of the specified incubation, the RNA was isolated using TRIzol reagent (Himedia), and cDNA was synthesized using iScript^TM^ cDNA synthesis kit (Biorad). qRT-PCR was performed using PowerUp™ SYBR™ Green Master Mix (Thermo Fisher Scientific). Relative expression of the gene was calculated using the 2^-ΔΔCt^ method by normalizing to reference gene *rpoD*. The primers used in RT-qPCR are listed in (Supplementary Table 2).

### In-silico analysis

The spacer sequences in CRISPR-I and CRISPR-II arrays were aligned with the coding and reverse complement sequences of different SPI-1 and SPI-2 genes using Serial Cloner version 2.6 software and screened for the presence of known protospacer adjacent motif (PAM) sequences in the vicinity.

### Statistical analysis

Statistical analysis was performed using Prism 8 software (GraphPad, California). Unpaired Student’s t-test was performed. Error bars indicate standard deviation (SD). Statistical significance is shown as follows: *p ≤ 0.05; ** p ≤ 0.01; *** p ≤0.001; ****,p < 0.0001; and ns, not significant.

## Results

### Deletion of the CRISPR-Cas system hampers invasion and intracellular survival of the CRISPR-Cas knockout strains in cell culture infection models

An essential feature of *Salmonella* pathogenesis relies on its ability to cross several physical and immunological defense barriers of the host invading a large variety of phagocytic and non-phagocytic cells. To understand the role of the CRISPR-Cas system in the pathogenicity of *S*. Typhimurium, we assessed the ability of the wildtype and CRISPR-Cas knockout strains to invade and proliferate in intestinal epithelial cells and macrophages. We evaluated the invasion and intracellular survival of the strains in non-polarized and polarized HT-29 cells, a colon carcinoma intestinal epithelial cell line. Compared to the wildtype strain, the knockout strains were impaired in invading the HT-29, and the invasiveness was rescued upon complementation with the knocked-out gene. However, the CRISPR-Cas knockout strains exhibited an enhanced attenuation in percentage invasion in polarized cells (Fig. 1A). Though the fold replication of all the strains was less in the polarized cells, the knockout strains showed reduced replication than that of the wildtype in both the cell types. The reduction in fold replication of the knockout strains was 1.5-2 times in non-polarized HT-29, while in polarized cells, the fold replication of knockout strains was ∼1.3 times less than that of the wildtype strain (Fig. 1B).

**Fig. 1:**
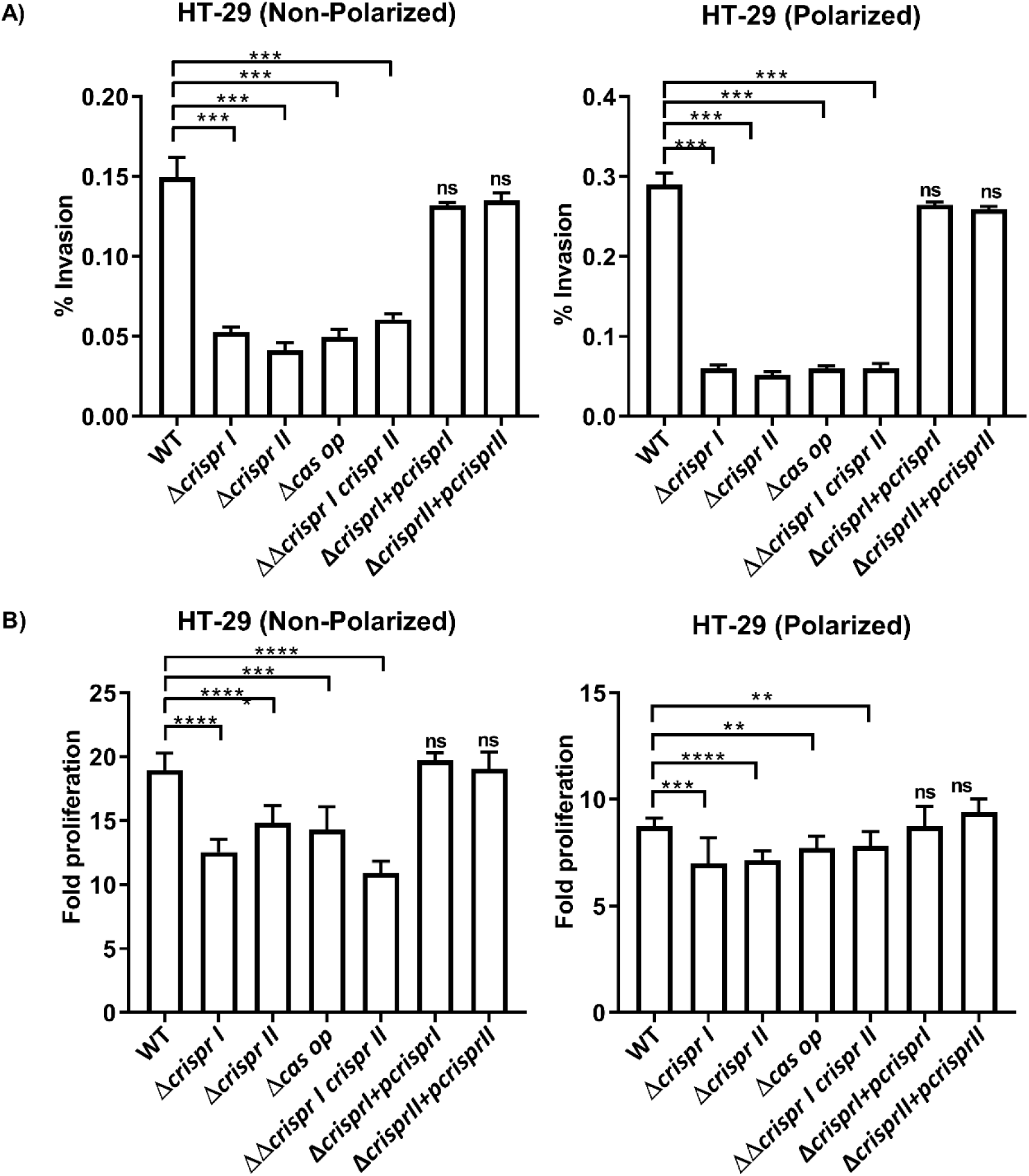
CRISPR-Cas knockout strains show invasion and replication defects in intestinal epithelial cells. **(A-B)** HT-29 cell lines, both non-polarized and polarized, were infected with *S.* Typhimurium strain 14028s wildtype (WT), CRISPR (Δ*crisprI,* Δ*crisprII,* and ΔΔ*crisprI crisprII*) and *cas operon (*Δ*cas op*) knockout strains along with their respective complements (Δ*crisprI+*p*crisprI* and Δ*crisprII+*p*crisprII*). **(A)** The percentage of invasion in intestinal epithelial cells was calculated using CFU analysis of the infected cell lysate and the pre-inocula used for infection. Fold proliferation was calculated by normalizing the CFU at 16 h to 2h. An student t-test was used to determine significant differences between the WT and knockout strains. Error bars indicate SD. Statistical significance is shown as follows: *, p ≤ 0.05; **, p ≤ 0.01; ***, p ≤0.001; ****, p < 0.0001; and ns, not significant.

After evading the intestinal epithelial barrier, *Salmonella* utilizes macrophages for its systemic dissemination. Therefore, we checked the invasion and intracellular survival of the CRISPR-Cas knockout strains in RAW 264.7 macrophage cell lines and peritoneal macrophages (considered immune sentinels with enhanced phagocytic activity). The percentage phagocytosis of the knockout strains by the macrophages was reduced to ∼ 35-60% of the phagocytosis of the wildtype strain (Fig. 2A). Moreover, the intracellular proliferation of the CRISPR-Cas knockout strains decreased by 1.5-2.5-fold in RAW 264.7 cell lines and by 2.5- 5-fold in peritoneal macrophages (Fig. 2B). In all these infection experiments, the complementation of corresponding genes in Δ*crisprI* and Δ*crisprII* showed results comparable to that of the wildtype, confirming that the gene deletion process did not produce any polar effects.

**Fig. 2:**
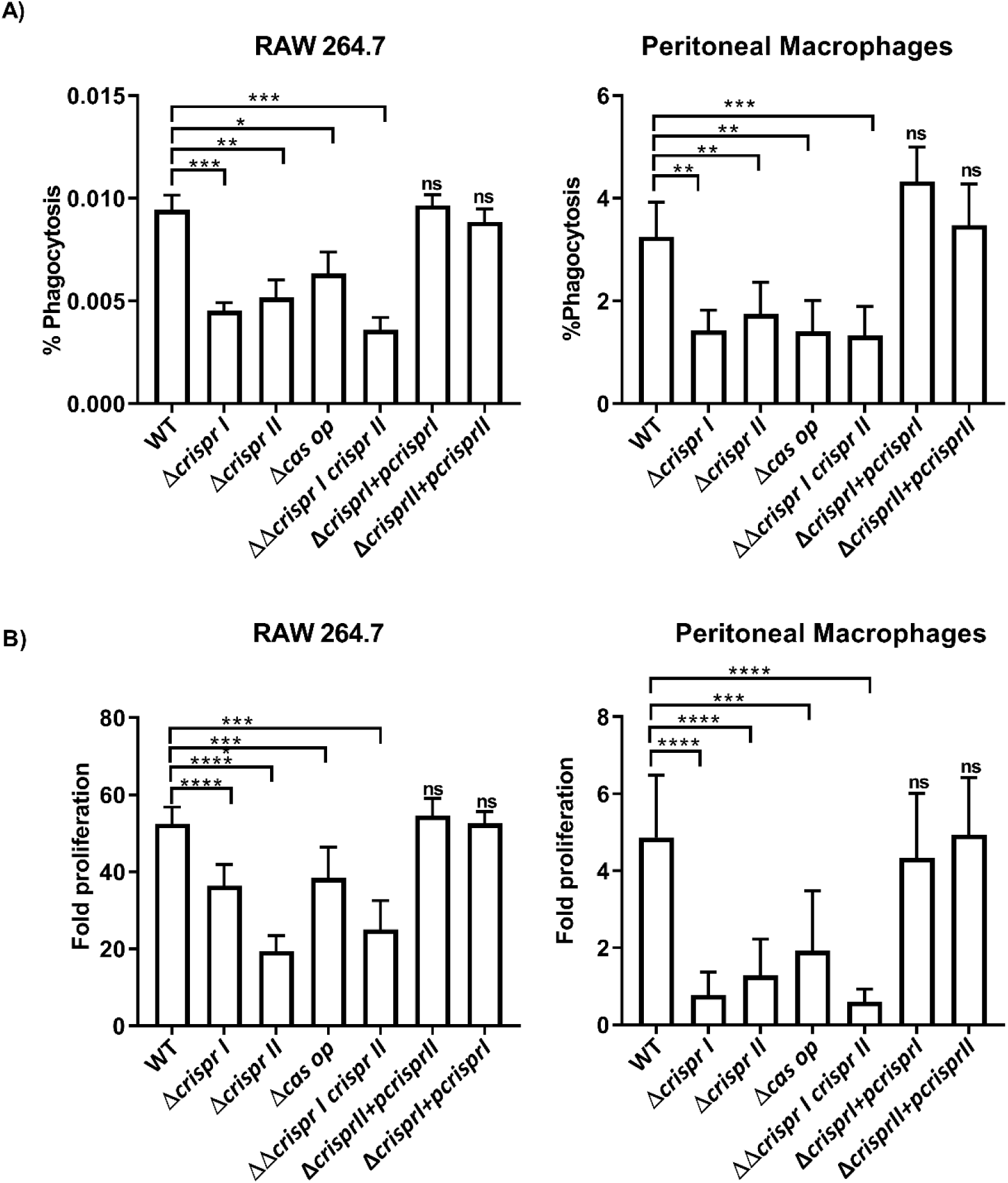
CRISPR-Cas knockout strains are attenuated in intracellular survival in macrophages. RAW 264.7 macrophage cell lines, and peritoneal macrophages were infected with *S.* Typhimurium strain 14028s wildtype (WT), CRISPR (Δ*crisprI,* Δ*crisprII,* and ΔΔ*crisprI crisprII)* and *cas operon (*Δ*cas op*) knockout strains along with their respective complements (Δ*crisprI+*p*crisprI* and Δ*crisprII+*p*crisprII*). **(A)** The percentage of phagocytosis by the macrophages was calculated using CFU analysis of the infected cell lysate and the pre-inocula used for infection. **(B)** Fold proliferation was calculated by normalizing the CFU at 16 h to 2h. An unpaired t-test was used to determine significant differences between the WT and knockout strains. Error bars indicate SD. Statistical significance is shown as follows:: *, p ≤ 0.05; **, p ≤ 0.01; ***, p ≤0.001; ****, p < 0.0001; and ns, not significant.

### Deletion of the CRISPR-Cas system attenuated virulence of *Salmonella* in *in-vivo* infection models

The impaired intracellular survival of the CRISPR-Cas knockout strains in different cell-culture infection models prompted us to look into the pathogenic potential of the CRISPR-Cas knockout strains in *in-vivo* infection models. Off-late, *C. elegans* has emerged as an attractive model host to study *Salmonella* pathogenesis. The pathogenic potential of the knockout strains was tested in *C. elegans* using the bacterial colonization assay. We observed a 40-60% reduction in the colonization of nematodes exposed to the CRISPR-Cas knockout strains compared to those exposed to the wildtype strain (Supplementary Figure S1). These observations were further validated in the murine model of typhoid fever using BALB/c mice. The infected mice were dissected three days post-infection to enumerate the bacterial burden in the Peyer’s patch (PP), mesenteric lymph node (MLN), liver, and spleen. The knockout strains displayed significantly reduced bacterial load in all these organs (Fig. 3A), indicating the role of the CRISPR-Cas system in establishing *in vivo* infection of *Salmonella*.

**Fig. 3:**
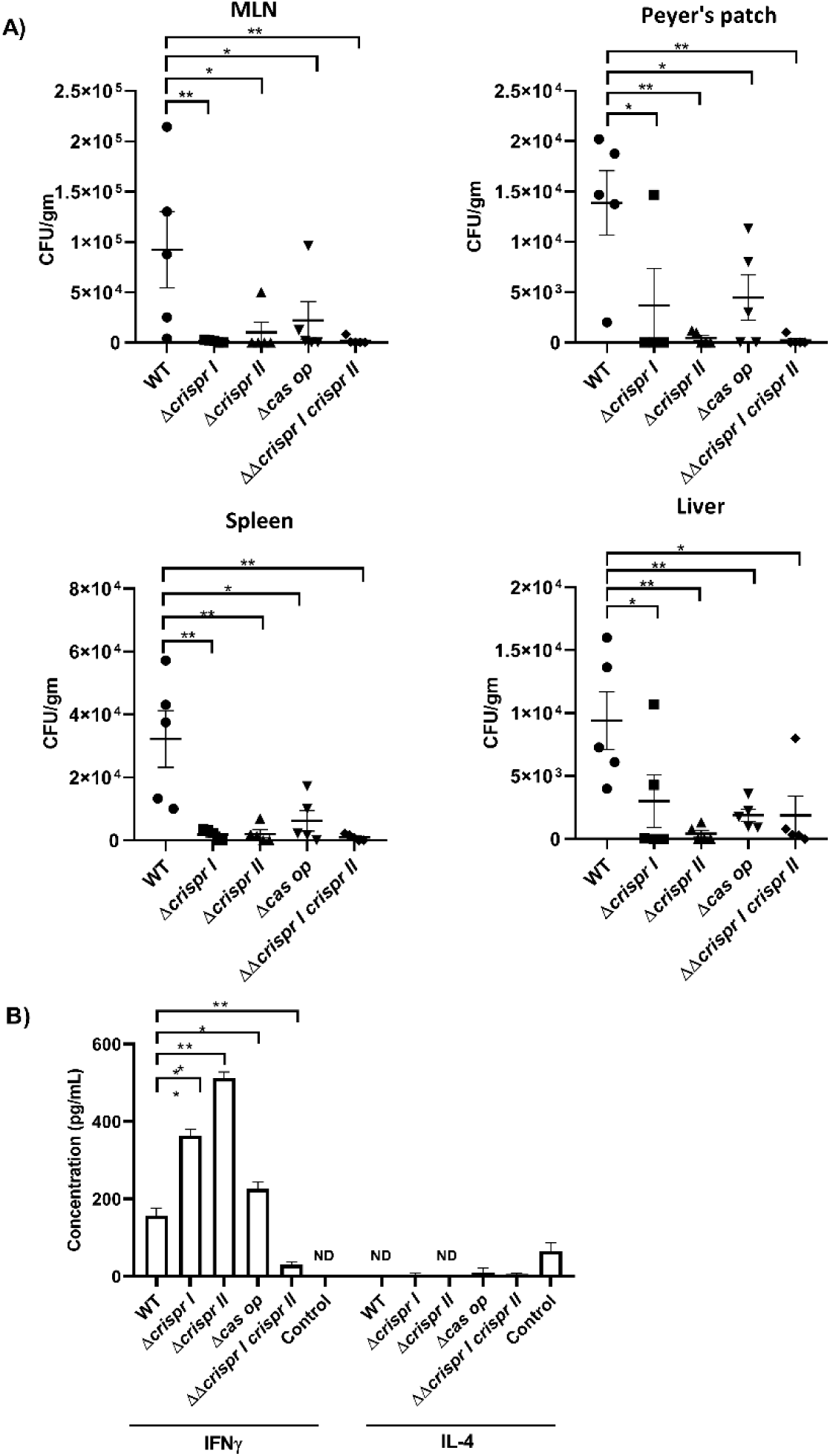
The CRISPR-Cas knockout strains show impaired colonisation in *in-vivo* model organism (mice) **(A)** Bacterial burden in different reticuloendothelial organs of the mice infected with wildtype (WT) and CRISPR (Δ*crisprI,* Δ*crisprII,* and ΔΔ*crisprI crisprII)* and *cas operon (*Δ*cas op*) knockout strains was estimated 3 days post-infection by CFU analysis. **(B)** The concentrations of proinflammatory cytokine IFN-γ, and anti-inflammatory cytokine IL-4 in the pooled sera of infected mice were determined using ELISA. Results are represented as mean ± SD pooled sera samples (2 mice per pool) for each infected and control group. An unpaired t-test was used to determine significant differences between the WT and knockout strains. Error bars indicate SD. Statistical significance is shown as follows: *, p ≤ 0.05; **, p ≤ 0.01; ***, p ≤0.001; ****, p < 0.0001; and ns, not significant.

Next, we assessed the inflammatory immune responses in the sera of the infected BALB/c mice. The pro-inflammatory cytokine, IFN-γ showed elevated levels in serum during infection, except that of ΔΔ*crisprI crisprII.* The anti-inflammatory cytokine IL-4 exhibited very low serum concentrations in mice infected with strains Δ*crisprI*, Δ*cas op*, and ΔΔ*crisprI crisprII*, in contrast to its absence in the serum of mice infected with the wildtype strain (Fig. 3B). Conversely, IL-10 was not detectable in the serum of the infected mice.

### The CRISPR-Cas knockout strains are susceptible to antimicrobial peptides, and the complement system

The intestinal invasion by *Salmonella* evokes innate immune responses by the host. The intestinal epithelial cells reinforce the intestinal barrier function by releasing antimicrobial peptides (AMPs), while the immunity components in serum, like lysozyme and complement (also present in the intestine), restrict microbial colonization [23]. *Salmonella* shows resistance against these antimicrobials by elongating or modifying the LPS O-antigen [24,25]. As the CRISPR-Cas knockout strains are attenuated in virulence, we estimated the sensitivity of the knockout strains against serum (complement system) and cationic AMPs like protamine sulphate and polymyxin B. Compared to the wildtype strain, the knockout strains showed ∼40-50% and ∼60-70% reduction in their percentage survival in the presence of protamine sulphate and polymyxin B, respectively (Fig. 4A). The survival of the knockout strains was reduced by ∼15-30 % in the presence of serum (Fig. 4A).

**Fig. 4:**
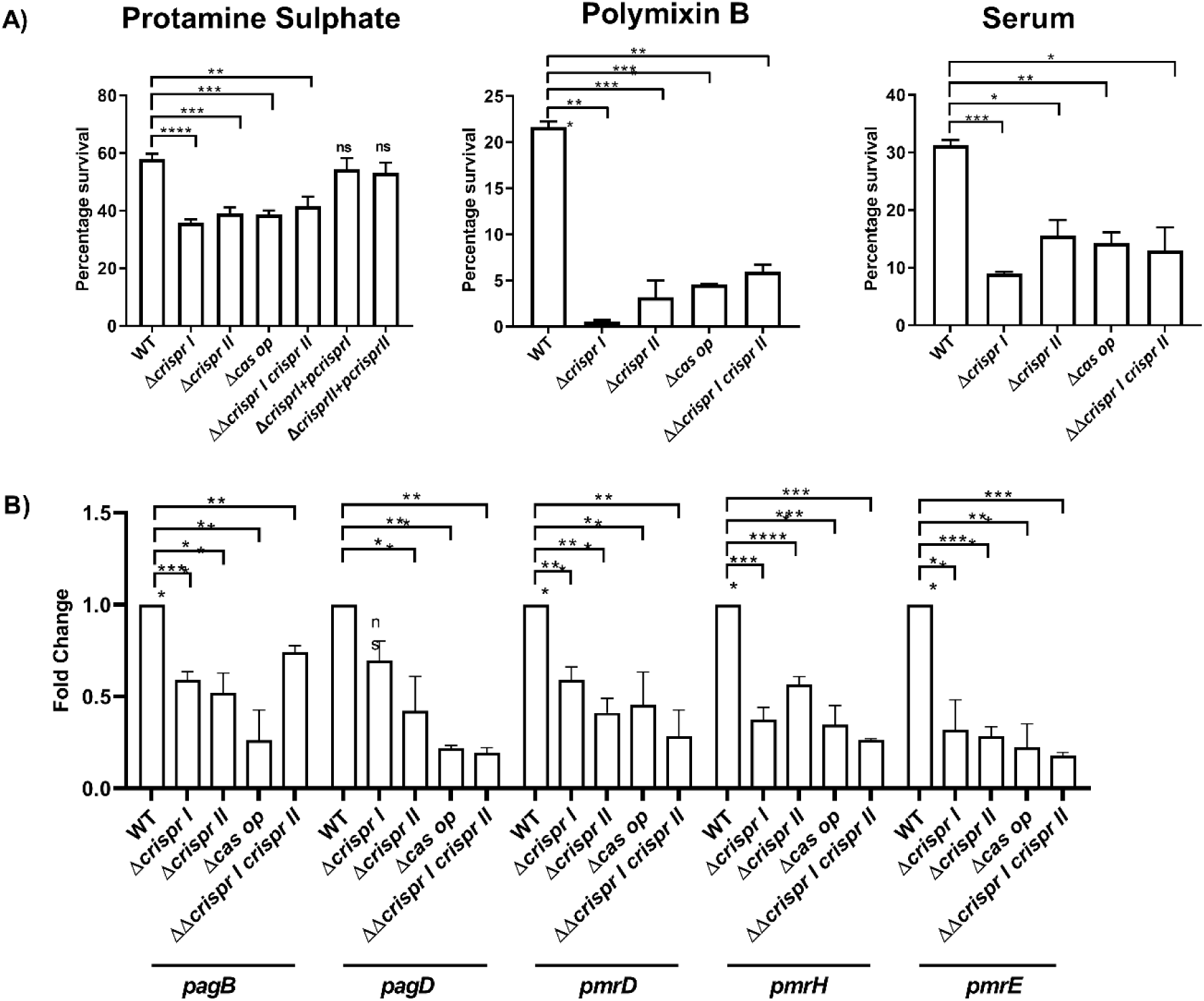
CRISPR-Cas knockout strains show sensitivity towards antimicrobial peptides (AMP), and the complement system. **(A)** AMP sensitivity of the mutant strains against the cationic peptides-protamine sulfate (0.5 μg/mL), polymyxin B (0.5 μg/mL), and serum (20% FBS) was tested by survival assay. The strains were exposed to antimicrobials and percentage survival was determined by analyzing CFU in both untreated and treated samples, with the untreated samples serving as controls. Percentage survival in serum was determined by using heat-inactivated samples as control. An unpaired t-test was used to determine significant differences between the WT and knockout strains. Error bars indicate SD. Statistical significance is shown as follows: *, p ≤ 0.05; **, p ≤ 0.01; ***, p ≤0.001; ****, p < 0.0001; and ns, not significant. **(B)** Total RNA isolated from late log-phase bacteria strains was used for cDNA synthesis, followed by qRT-PCR to assess the expression of polymyxin resistance (*pmr*) genes like *pagB*, *pagD*, *pmrH*, *pmrE*, *pmrD.* Relative expression of the gene was calculated using the 2 ^-ΔΔCt^ method, and normalized to reference gene *rpoD*.

Gram-negative bacteria modify their lipopolysaccharides as a strategy of protection against substances like polymyxins[26]. These modifications include replacing phosphate groups with L-Ara4N, or PEtN, thus reducing the negative charge of lipid A, thereby hindering the AMP binding and leading to AMP resistance[27]. Two-component systems, like PhoP/PhoQ and PmrA/PmrB, trigger the upregulation of operons like *pmrCAB* and *arnBCADTEF-pmrE* (*pmrHFIJKLM-ugd*), facilitating the synthesis and transfer of PEtN and L-Ara4N to lipid A [27]. Therefore, we examined the expression patterns of several polymyxin resistance *(pmr*) genes, namely *pagB*, *pagD*, *pmrH*, *pmrE*, *pmrA* and *pmrD.* The knockout strains exhibited a 1.4-to-2.5-fold decrease in expression levels of *pagB*. Similarly, the transcripts of *pagD* (∼1.4 to 2.5 fold)*, pmrD* (∼2 to 2.5 fold), *pmrH* (∼2.5 to 3 fold) and *pmrE* (2.5-fold) were downregulated in all the knockout strains (Fig. 4B). However, there was no consistent change in the expression of the *pmrA* gene across all the strains (Supplementary Figure S2).

Collectively the results indicate that the CRISPR-Cas knockout strains have an impaired ability to overcome innate immune barriers during the dissemination and intestinal infection phase.

### The CRISPR-Cas knockout strains show altered expression of *Salmonella* pathogenicity island (SPI-1 and SPI-2) genes

Alterations in the expression of *Salmonella* pathogenicity islands (SPI-1 and SPI-2) genes could be one of the contributing factors to diminished host cell invasion and intracellular replication by the CRISPR-Cas knockout strains. Thus, to gain mechanistic insights into the regulation of pathogenesis by the CRISPR-Cas knockout strains, we checked the expression of effectors encoded by SPI-1 and SPI-2 pathogenicity island using RT-PCR.

The SPI-1, required during the intestinal phase of infection, delivers the effector proteins required for intestinal invasion and inflammation inside the host cells [28]. As the CRISPR and Cas knockout strains were defective in the invasion, we first assessed the expression of SPI-1 regulatory genes like *hilD, hilA,* and *h-ns*. All the knockout strains showed reduced expression (∼3-4 fold) of *hilA,* whereas the *hilD* transcript was significantly downregulated by just 1.2 to 1.7∼fold in all the knockout strains (Fig. 5A). The *h-ns* gene did not show any difference in the expression among all the strains (Supplementary Figure S3). Next, to envisage the impaired invasion ability of the knockout strains, we analysed the expression of a few important SPI-1 effectors, *sipA, sipD,* and *sopB*. The CRISPR-Cas knockout strains showed ∼2-2.5 fold and ∼1.5-2 fold reduced expression of *sipA* and *sipD,* respectively. However, *sopB* was downregulated by only ∼ 1.4-fold in the knockout strains, except for ΔΔ*crisprI crisprII,* which showed more than two-fold downregulation (Fig. 5A).

**Fig. 5:**
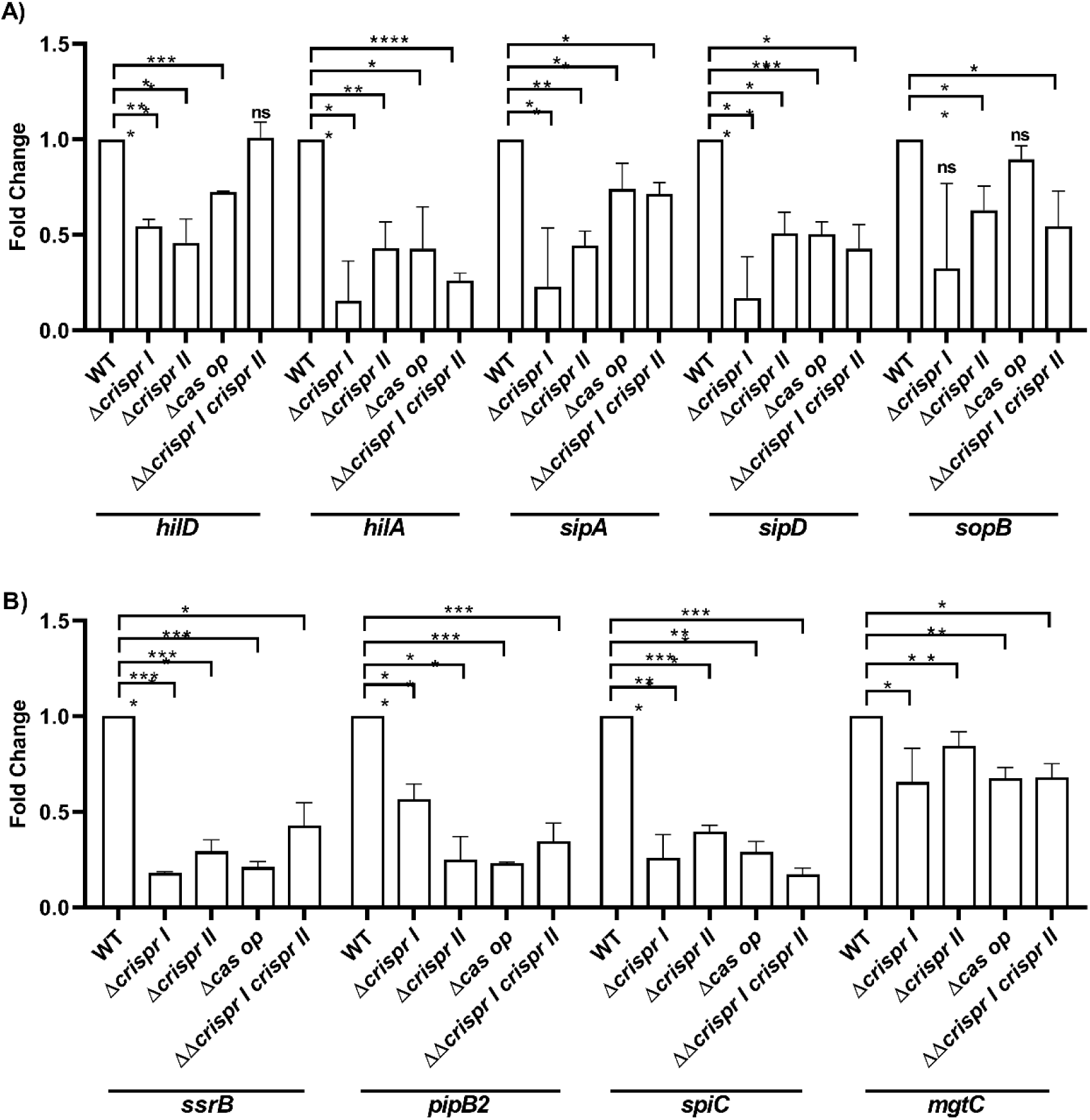
The CRISPR-Cas system regulates SPI-1, and SPI-2 genes expression. **(A)** The bacterial strains were cultivated in conditions that promote SPI-1 activation. Subsequently, qRT-PCR was conducted on isolated RNA samples to assess the expression levels of key SPI-1 components, including transcriptional activator-*hilD*, transcriptional regulator-*hilA*, and SPI-1 effectors- *sipA*, *sipD*, *sopB*. **(B)** The bacterial strains were cultivated in SPI-2 inducing conditions for 5 h, and qRT-PCR was performed from isolated RNA to check expression of SPI-2 effectors- *pipB2* and *spiC*, SPI-2 encoded transcriptional regulator-*ssrB*, and SPI-3 encoded protein-*mgtC*. Relative expression of the gene was calculated using the 2 ^-ΔΔCt^ method and normalized to the reference gene, *rpoD*.

Following the epithelial cell invasion, *Salmonella* employs SPI-2 encoded effector proteins to form a permissive-replicative niche in SCV [29]. SPI-2 expression is induced within SCV and is controlled by a two-component system, SsrAB [30]. The SsrB drives the expression of various SPI-2 effector proteins like PipB2, SpiC, etc. The expression of representative SPI-2 genes and its regulator SsrAB was checked in the strains grown in F-media. The expression of the SPI-2 effector, *pipB2* and *spiC*, and the transcriptional regulator, *ssrB,* was downregulated by more than 2-fold in all the knockout strains (Fig. 5B). The low Mg^2+^ milieu of SCV promotes MgtC expression, a virulence protein required for intracellular replication inside macrophages [31]. Hence, we also evaluated the expression of *mgtC* in strains grown in F-media. All the knockout strains show 1.25-1.6-fold downregulation in *mgtC* expression (Fig. 5B).

### The CRISPR-Cas knockout strains show reduced survival against oxidative response but induce similar oxidative responses in macrophages as that of the wildtype

During the early and late stages of infection, *Salmonella* encounters oxidative and nitrosative stress inside macrophages, as a part of the host’s immunological response [32]. The diminished intracellular proliferation of the CRISPR-Cas knockout strains in macrophages may be attributed to their ability to induce elevated production of free radicals, leading to enhanced cellular death. Alternative explanations for the reduced survival include the strains’ increased susceptibility to oxidative and nitrosative stress responses triggered by the host cells.

At first, we assessed the viability of the knockout strains in the presence of oxidative stressors like H_2_O_2_ and nitric oxide. In the presence of H_2_O_2_, the percentage survival of the knockout strains was significantly reduced by ∼60-70% for Δ*crisprI*, Δ*crisprII*, Δ*cas op*, and ΔΔ*crisprI crisprII* when compared to that of the wildtype (Fig. 6A). However, all the strains exhibited similar sensitivity to sodium nitrite (Supplementary Figure S4).

**Fig. 6:**
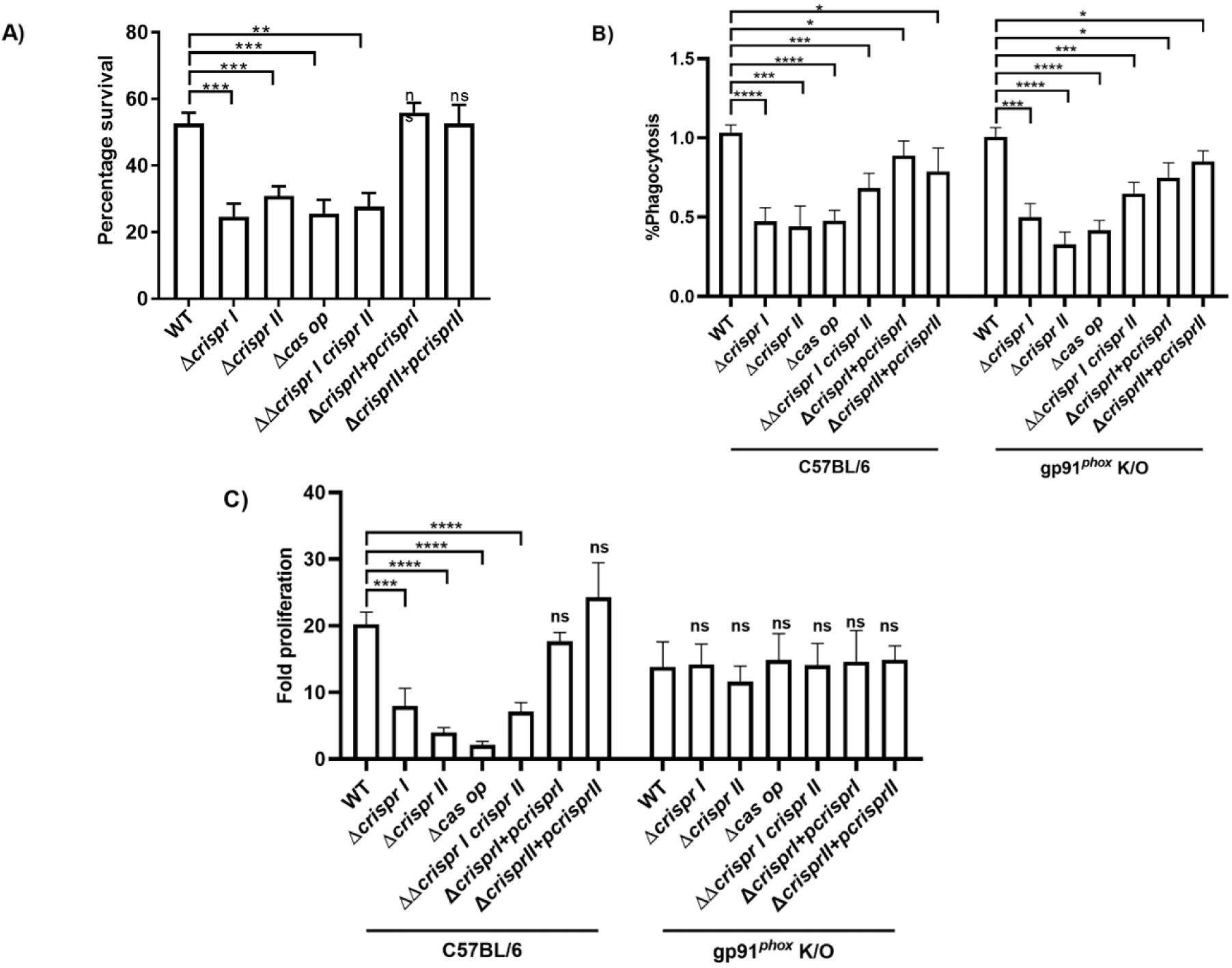
CRISPR-Cas knockout strains’ diminished proliferation in phagocytic cells could be linked to their susceptibility to ROS. **(A)** Overnight-grown strains were exposed to 1 mM H_2_O_2_ for 2 hours at 37°C in the dark. CFU counts were then determined, and percentage survival was calculated by normalizing treated samples to untreated controls. **(B)** The percentage of phagocytosis by peritoneal macrophages derived from GP91*^phox^* knockout mice was determined through CFU analysis of infected cell lysates. Intracellular fold proliferation was calculated by normalizing the CFU count at 16 h to 2h. An unpaired t-test was used to determine significant differences between the WT and knockout strains. Error bars indicate SD. Statistical significance is shown as follows: *, p ≤ 0.05; **, p ≤ 0.01; ***, p ≤0.001; ****, p < 0.0001; and ns, not significant.

Following this, the strains’ intracellular survival was assessed in macrophages with enhanced (LPS induced) and suppressed (gp91*^phox^* knockout) oxidative response. Similar to the data with the wildtype peritoneal macrophages, the percentage phagocytosis of the CRISPR-Cas knockout strains by the gp91*^phox-/-^* macrophages was reduced to ∼50-60% that of the wildtype *Salmonella* (Fig. 6B). As anticipated, the intracellular proliferation of the CRISPR-Cas knockout strains was comparable to that of the wildtype (Fig. 6B). Next, we assessed the proliferation of these strains in macrophages with enhanced and suppressed ROS production (Supplementary Figure S5A). The CRISPR-Cas knockout strains exhibited comparable intracellular proliferation to that of wildtype *Salmonella* in conditions of suppressed oxidative response (Supplementary Figure S5C).

When assessed for induction of oxidative response in the macrophages by the CRISPR-Cas knockout strains, we found comparable intracellular ROS levels in macrophages infected with the knockout or wildtype strains (Supplementary Figure S5A). Likewise, no significant difference was observed for the extracellular nitric oxide produced by these infected cells (Supplementary Figure S5B).

### CRISPR-Cas knockout strains exhibit susceptibility to ROS as a result of amplified H_2_O_2_ influx and diminished expression of antioxidant genes

It is known that the outer membrane porin, OmpW aids the influx of H_2_O_2_ in *Salmonella* [33], and the *ompW* null mutants of *E. coli* and *Salmonella* are resistant to oxidative stress [33,34]. Additionally, the *ompW* null mutants show enhanced surface-attached biofilm in *Cronobacter sakazakii*[35,36]. The CRISPR-Cas knockout strains showed reduced H_2_O_2_ tolerance (Fig. 6A) and lesser surface-attached biofilm compared to that of wildtype[16]. We hypothesized that the upregulated *ompW* in the CRISPR-Cas knockout strains might be leading to the observed phenotypes. With this antecedent, we assessed the expression of *ompW* in the knockout strains. The *ompW* expression was 2-fold higher for Δ*crisprI* and Δ*cas op*, while in Δ*crisprII* and ΔΔ*crisprI crisprII* it was 3-fold higher than that of the wildtype (Fig. 7A). H_2_O_2_ is known to repress the expression of *ompW* in *S*. Typhimurium [33]. As expected, the H_2_O_2_ treatment reduced the expression of *ompW*, and the difference in the expression between the knockout strains and wildtype reduced to 1 to 1.3-fold. Considering that the *ompW* is expressed at similar levels in H_2_O_2_ primed wildtype and knockout strains, we next assessed the survival of the knockout strains post-priming. The percentage survival of the primed knockout strains was similar to that of the primed wildtype (Supplementary Figure S6).

**Fig. 7:**
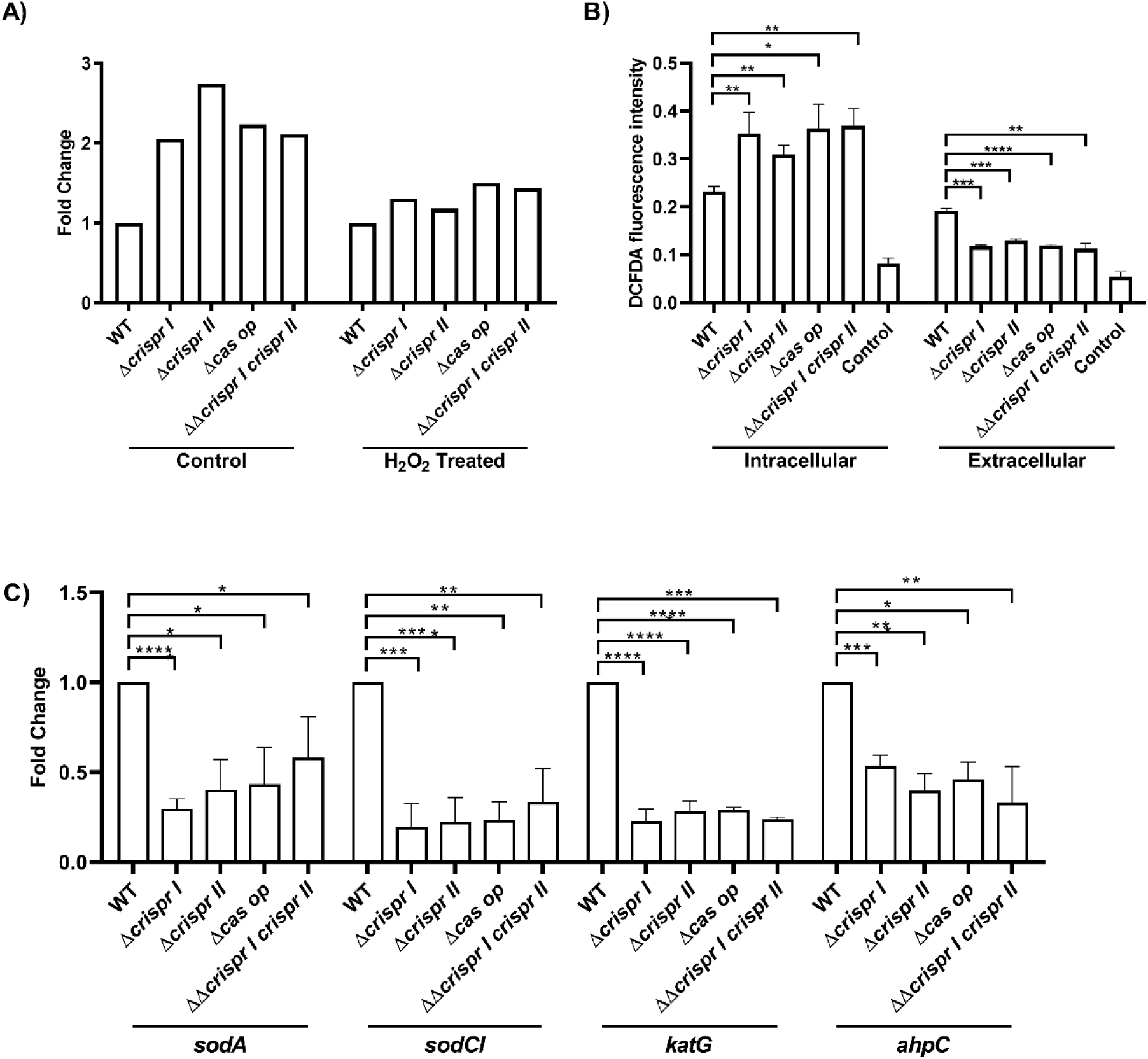
The CRISPR-Cas knockout strains are susceptible to ROS owing to elevated influx H_2_O_2_, and reduced expression of antioxidant genes. **(A)** The bacterial strains were cultivated in F-media, with and without H_2_O_2_ for 8 h, followed by qRT-PCR analysis of *ompW*. Relative expression of the gene was calculated using the 2 ^-ΔΔCt^ method and normalized to reference gene *rpoD*. **(B)** The bacterial strains were cultivated in F-media until they reached an OD_600nm_∼0.5. They were then incubated in the dark for 5 minutes with 1 mM H_2_O_2_. The H_2_O_2_ levels in both extracellular and intracellular fractions were measured using H_2_DCFDA. The H_2_O_2_ untreated sample was used as a control. **(C)** The bacterial strains were cultivated in SPI-2 inducing conditions, and qRT-PCR was performed from isolated RNA to evaluate the expression of ROS detoxifying enzymes, superoxide dismutases (*sodCI* and *sodA*), catalase (*katG*), and peroxidase (*ahpC*). Relative expression of the gene was calculated using the 2 ^-ΔΔCt^ method and normalized to the reference gene, *rpoD.* An unpaired t-test was used to determine significant differences between the WT and knockout strains. Error bars indicate SD. Statistical significance is shown as follows: *, p ≤ 0.05; **, p ≤ 0.01; ***, p ≤0.001; ****, p < 0.0001; and ns, not significant.

To combat the oxidative stress response generated by the host cell*, Salmonella* employs an array of antioxidant enzymes like superoxide dismutase, catalase, and peroxidase to detoxify ROS [37]. As the knockout strains are sensitive to H_2_O_2_, we analyzed the expression of antioxidant genes (*sodA, sodCI, katG,* and *ahpC)* in the strains grown in F-media. The antioxidant genes were repressed in all the knockout strains (Fig. 7C) explaining their reduced survival in the presence of H_2_O_2_.

### Partial binding of crRNA to genes may lead to the regulation of certain genes

The CRISPR spacers, specifically spacers 2 and 8 from the CRISPR-I array and spacer 18 of CRISPR-I array, demonstrated partial complementarity (46.875% to 56.25%) with SPI-1 effectors *sopB* and *sipA*, respectively. Additionally, spacer 7 of the CRISPR-II array showed partial complementarity (56.25% to 59.375%) with *hilA*.

For SPI-2, the transcriptional regulator *ssrB* and the effector *pipB2* displayed partial complementarity with spacer 22 of CRISPR-I (43.75% to 46.875%) and spacer 20 of CRISPR-II (53.125%), respectively. Furthermore, other genes such as superoxide dismutases (*sodA*), catalase (*katG*), and outer membrane porin (*ompW*) exhibited partial complementarity with spacers 4 and 23 of the CRISPR-I array (46.875% to 53.125%), spacers 11 of the CRISPR-I array (43.75% to 53.125%), and spacers 3 of the CRISPR-I array ( 53.125%), respectively.

In summary, independent regulation of SPI-1 and SPI-2 genes, and antioxidant genes by the CRISPR-Cas system may occur due to a partial match between the CRISPR spacers and the genes (Supplementary Figure S7).

## Discussion

Our present study demonstrates that the CRISPR-Cas knockout strains of *S*. Typhimurium are attenuated in virulence, showing reduced invasion and proliferation in host cells.

*Salmonella* invasion of non-phagocytic cells is mediated by SPI-1-encoded effectors (Fig. 8) [38]. The SPI-1 transcriptional regulator, *hilD*, binds upstream of the master regulator, *hilA*, thereby counteracting its *h-ns*-mediated repression[19]. HilA in turn regulates the expression of the T3SS’s structural and effector proteins like Sips and Sops[19]. The reduced expression of *hilA* in the CRISPR-Cas knockout strains may explain the downregulation of the other T3SS structural and effector proteins (Fig. 8).

**Fig. 8:**
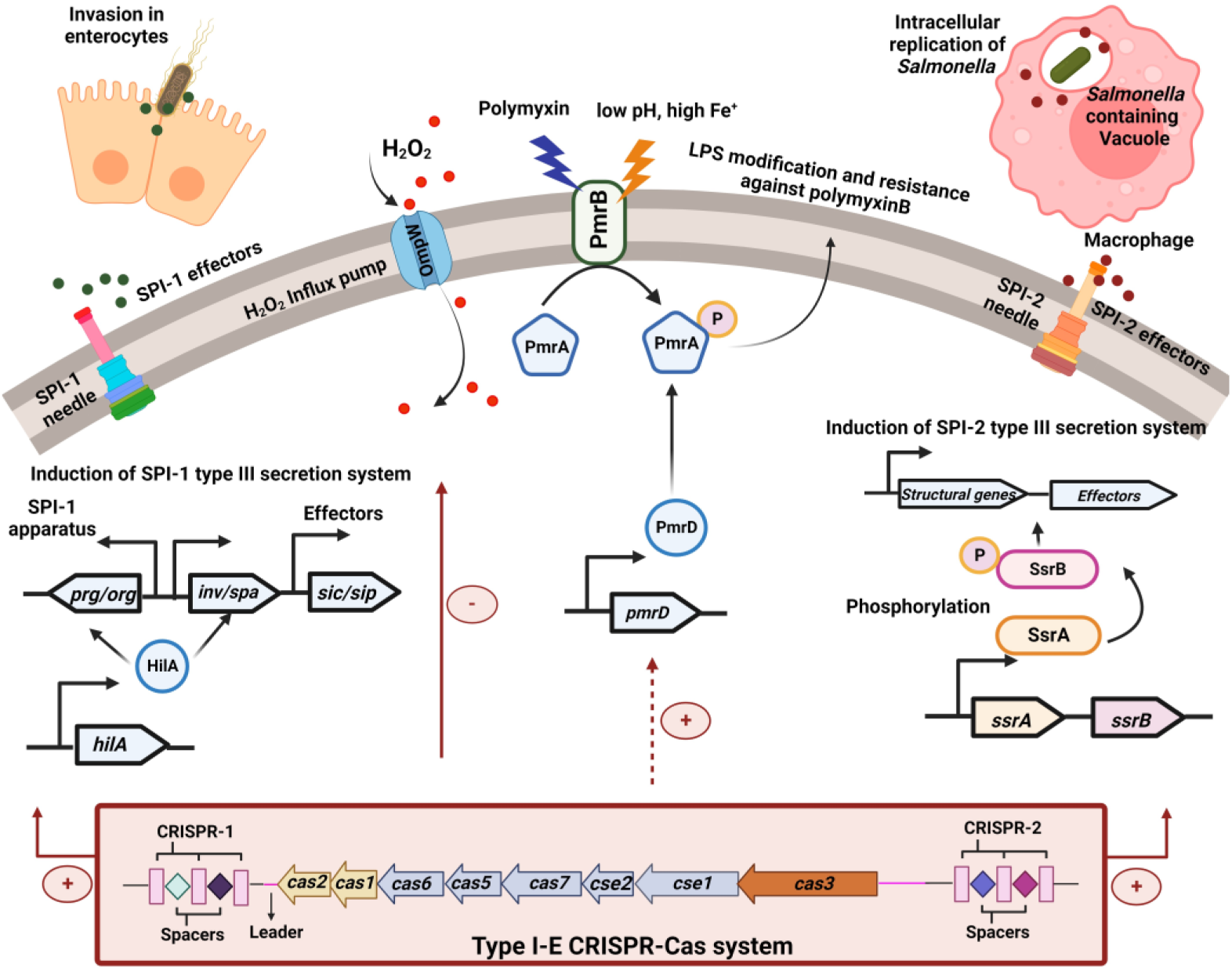
Deletion of the CRISPR-Cas system attenuates *Salmonella* pathogenicity. Proposed mechanisms of the type I-E CRISPR-Cas system in regulating *Salmonella* pathogenesis via modulation of SPI-1 and SPI-2 genes. We propose that the CRISPR-Cas system positively regulates *hilA* (direct regulation via complementary base-pairing between crRNA and gene) whereby it upregulates the expression of SPI-1 apparatus and effector proteins (direct regulation of *sipA*) involved in the invasion of enterocytes by *Salmonella*. The intestinal epithelial cells reinforce the intestinal barrier function by releasing antimicrobial peptides. The CRISPR-Cas system appears indirectly regulate (red dotted lines) *pmr* genes to provide resistance against antimicrobial peptides (AMPs). Within the *Salmonella* containing vacuole (SCV) of macrophages, the bacteria shut down its SPI-1 system and activates the SPI-2-encoded SsrAB system in response to the acidic milieu. The SsrAB system further activates the SPI-2 encoded genes. The CRISPR-Cas may be positively regulating SsrB (direct regulation) to trigger activation of SPI-2 encoded structural genes and effector proteins (direct regulation of *pipB2*) to aid intracellular proliferation and survival of *Salmonella*. In addition, the CRISRP-Cas system negatively regulates OmpW (direct regulation) during oxidative stress, thereby aiding in *Salmonella’*s survival. Taken together, the CRISPR-Cas system positively regulates *Salmonella* pathogenesis. The figure was created using Biorender.

SPI-1 translocases SipB, SipC, and SipD, are essential for the attachment of bacteria to the target cells [39], and SipA is required for the efficient invasion of *S*. Typhimurium during the early stages of infection [40]. Following membrane ruffling, *Salmonella* outer membrane proteins (Sops) control cytoskeletal rearrangement during the invasion and regulate polymorphonuclear leukocyte influx [41]. *In vivo* experiments demonstrate that SopB is required during the initial invasion process but also in the later stage of murine salmonellosis [42]. The knockout strains show decreased expression of the SPI-1 genes like *sipA*, *sipD,* and *sopB,* explaining their decreased invasion phenotype in *in-vitro* and *in-vivo* models. Partial complementarity between CRISPR spacers and *sipA* and *sopB* suggests the plausible involvement of the CRISPR-Cas system in modulating the regulation of these specific genes. Further, the altered LPS-O antigen structure significantly affects SipA translocation during invasion [43]. Thus, the decreased expression of *sipA* could also be attributed to the altered LPS structure of the CRISPR-Cas knockout strains [16].

Adhesins like flagella and Curli inherently contribute to adhesion and invasion into the epithelial cells [44,45]. Our previous studies demonstrated reduced expression of the flagellar and curli genes in the CRISPR-Cas knockout strains[16], thereby explaining their attenuated invasion in epithelial cells. To invade the epithelial cells, *Salmonella* relies on the flagellar subunit FljB to cross the mucosal barrier produced by the goblet cells [23]. The reduced invasion of the CRISPR-Cas knockout strains in differentiated HT-29 cells that produce mucin, over undifferentiated cells could be explained through the reduced *fljB* expression in these strains. This along with the increased sensitivity of the knockout strains to serum complement and AMPs present in the intestinal lumen, could explain their reduced colonization of the PP. It is reported that LPS modification affects bacterial susceptibility to complement [24], AMPs [46] and phagocytosis[24,43]. The reduced expression of LPS modifying genes along with *rfa a*nd *rfb* genes, coding for LPS core synthesis and O-antigen synthesis, respectively [16] in the knockout strains, may explain their defective phagocytosis and increased sensitivity to AMPs and complement.

Following bacterial uptake, the phagocytes elicit an oxidative burst. The toll-like receptors (TLRs), present on infected cells recognize *Salmonella*-derived ligands like LPS (TLR4), flagellin-FliC (TLR5), CsgA (TLR2), etc., thereby inducing a respiratory burst [23]. Though our data demonstrate altered expression of PAMPs in the CRISPR-Cas knockout strains [16], they fail to show any differences in the induction of oxidative response in RAW 264.7 cells compared to that of the wildtype. Despite comparable induction of ROS/RNS in the macrophages infected with the knockout and wildtype strains, the knockout strains have attenuated proliferation, possibly due to their increased susceptibility to H_2_O_2_ *via* the increased expression of OmpW, an importer of H_2_O_2_ (Fig. 8) [33]. The CRISPR-Cas system has also been reported to regulate outer membrane proteins in *S*. Typhi [47]. As the Omp protein is widely distributed in the Enterobacteriaceae family [48], the question is can the CRISPR-Cas system regulate *omp* expression in other members of the Enterobacteriaceae family? This question needs to be explored. OMPs and LPS help the bacteria tolerate different environmental stresses including, H_2_O_2_. *Salmonella* O-antigen capsule mutants are susceptible to H_2_O_2_ under biofilm conditions[49]. Thus, it may be possible that the altered LPS profile of the knockout strains also contributes to their H_2_O_2_ sensitivity. The sensitivity of the knockout strains to H_2_O_2_ corroborates the findings from our previous study, where a few cells in the CRISPR-Cas knockout strains become filamentous at 24 h [16]. This indicates a potential induction of reactive oxygen species (ROS) during biofilm formation and a few cells become filamentous in response to oxidative stress [16]. As the biofilm formation progressed, the nutrient deprivation could have accelerated ROS that could have been influxed in the knockout strains, resulting in their reduced viability at 24 h.

Apart from intracellular ROS, *Salmonella* also encounters extracellular H_2_O_2_ during the intestinal phase of infection [50], wherein it employs an array of oxidative enzymes to scavenge and degrade H_2_O_2_ molecules. Such enzymes include the cytoplasmic catalases (*katE, katG,* and *katN*), peroxidases (*ahpC*, *tpx*, and *tsaA*), superoxide dismutases (*sodA* and *sodB*), and the periplasmic superoxide dismutases (*sodCI*) [37]. The CRISPR-Cas knockout strains showed downregulation of these enzymes (one representative of each group), thereby displaying increased sensitivity against H_2_O_2_ and reduced survival within the macrophages and mice.

Besides the above-mentioned mechanisms, the coordinated action of other virulence determinants plays a major role in governing the survival and replication of *Salmonella* within SCV. Among them, MgtC is one such virulence factor that promotes intra-phagosomal replication under low Mg^2+^conditions [31]. It also promotes *Salmonella* virulence by negatively regulating cellulose production [51]. Deletion of *mgtC* attenuates *Salmonella* virulence in the mammalian host [51]. Interestingly, our study displayed such a relation wherein all the CRISPR-Cas knockout strains showed decreased *mgtC* expression and enhanced cellulose secretion [16]. Thus, supporting their impaired intracellular survival in phagocytic cells and *in vivo* models.

*Salmonella* is transported from the intestinal lumen and PP to the MLN, liver, and spleen either as extracellular bacteria or within the phagocytic cells [52]. Thus, the pathogen needs to overcome the innate immune barrier, like resistance against AMPs, ROS and serum proteins, to disperse systemically. The CRISPR-Cas knockout strains show reduced survival in the presence of AMPs, H_2_O_2,_ and serum, explaining their attenuated colonization and systemic spread. The differential expression of *pmr* genes (Fig. 8) together with altered LPS profile in the knockout strains could be one of the reasons for their increased sensitivity against AMPs and serum. However, the role of the CRISPR-Cas system in regulating other factors like OMPs, Rck and siderophore cannot be ruled out.

Following the internalization by phagocytic cells, the SPI-2-encoded SsrAB system gets activated in response to the acidic milieu. SsrA kinase phosphorylates the key regulatory factor of SPI-2, SsrB, which in turn activates the expression of SPI-2 encoded effector proteins like SpiC, PipB2, etc. (Fig. 8). Decreased expression of these and other virulence genes like *mgtC, katG, sod* and *ahpC* in the knockout strains could explain their sensitivity to antimicrobial defences, like ROS [53]. This could have have attenuated their virulence impacting colonization in *C. elegans* and mice. Furthermore, *in-silico* analysis shows partial complementarity between CRISPR spacers and key regulators (*ssrB*, *pipB2*)/other virulence-related genes (*sodA*, *katG*, *ompW*), suggesting potential regulatory interactions. Our findings underscore the involvement of the CRISPR-Cas system in governing SPI-1, SPI-2, and additional virulence genes crucial for *Salmonella’s* pathogenic behaviour (Fig. 8). However, further investigation is needed to unveil the specific regulatory mechanisms at play.

## Acknowledgements

We are very thankful to Jagannath Pradhan for his contribution to conducting experiments pertaining to real-time PCR. We would like to express our sincere gratitude to Birla Institute of Technology and Science, Pilani for their invaluable infrastructural support.

