## Supplementary figures and images for "Clustered Regularly Interspaced Short Palindromic Repeats-Cas system regulates *Salmonella* virulence"

### Supplementary Figures

#### Slide 1
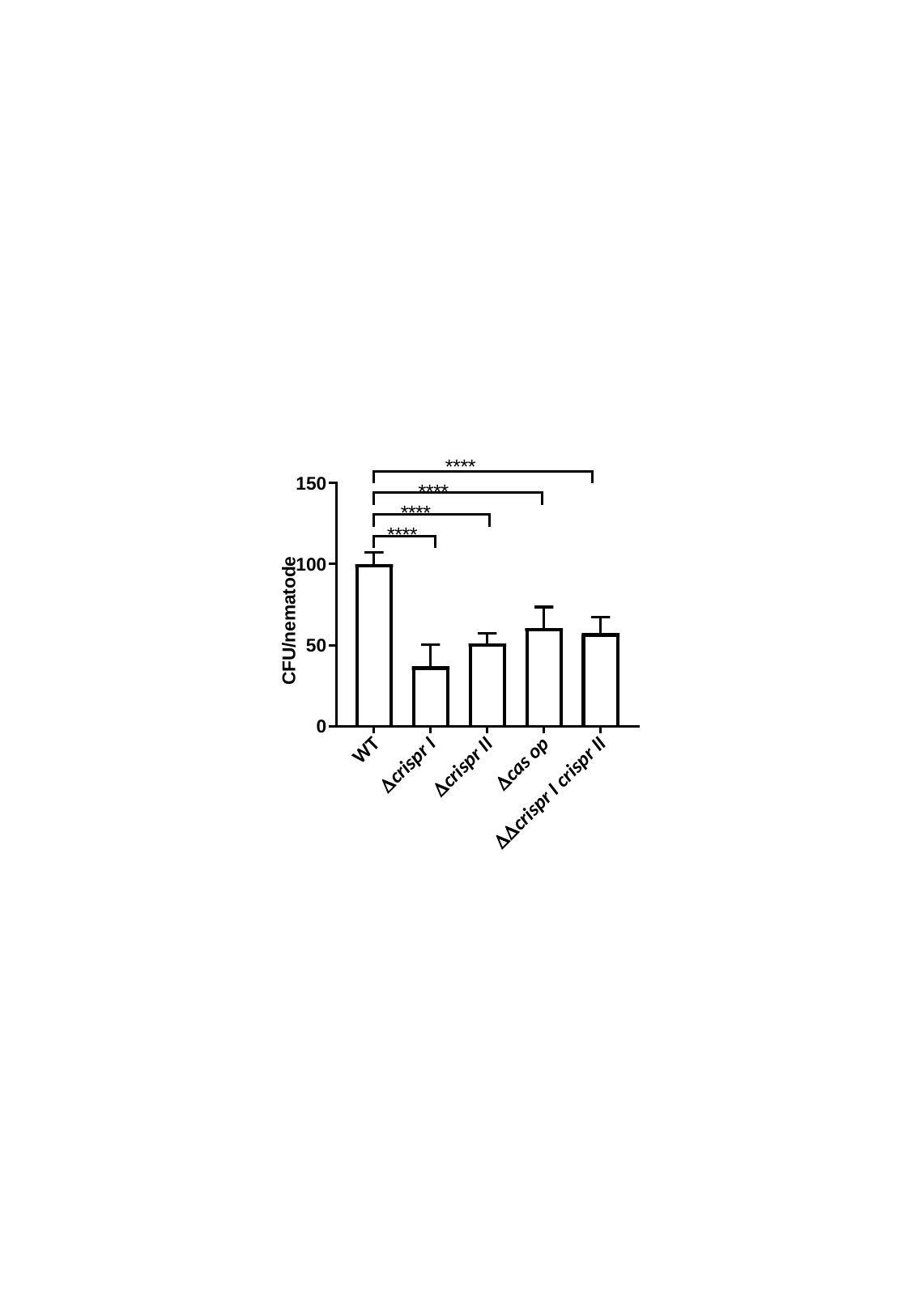

#### Slide 2
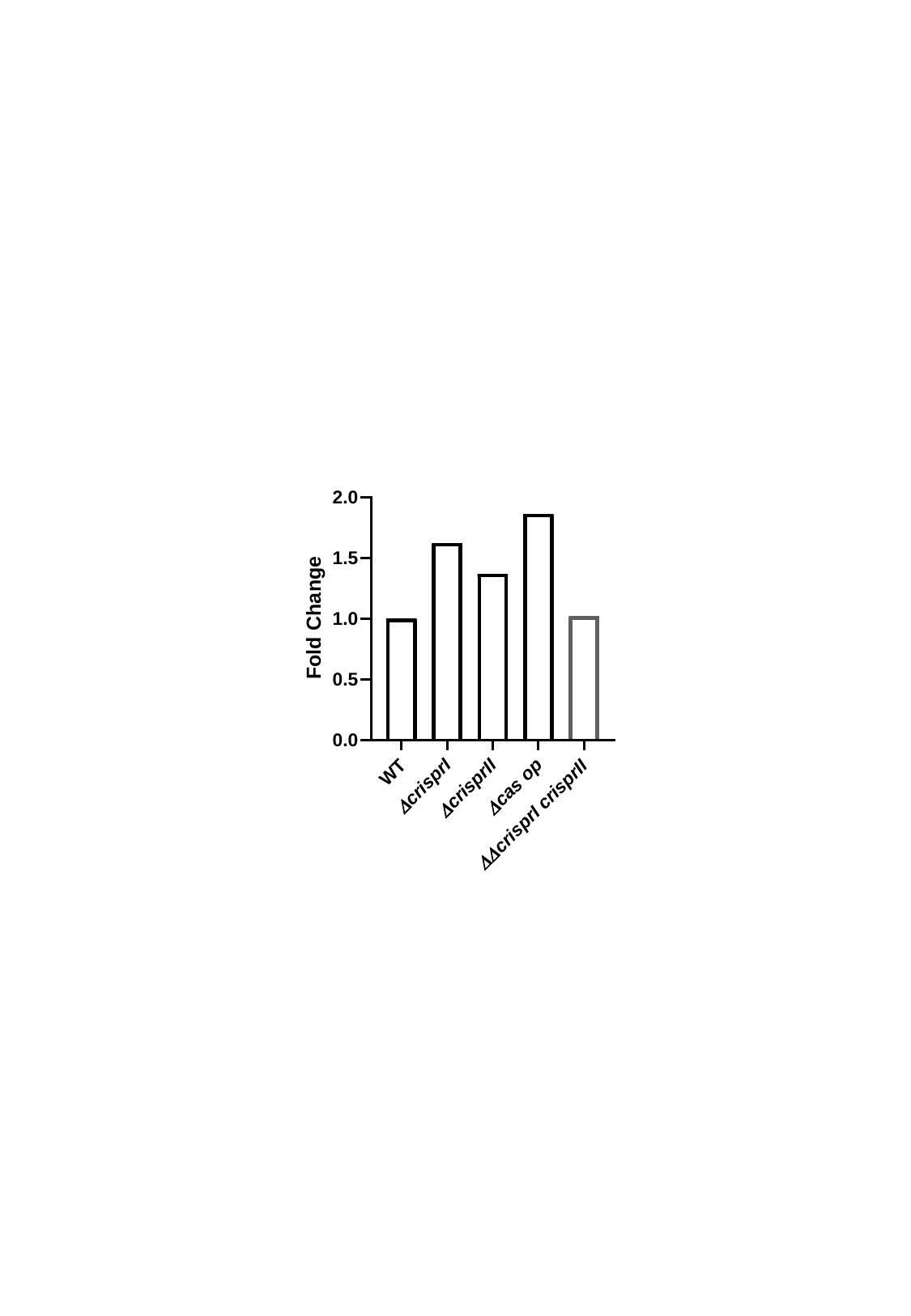

#### Slide 3
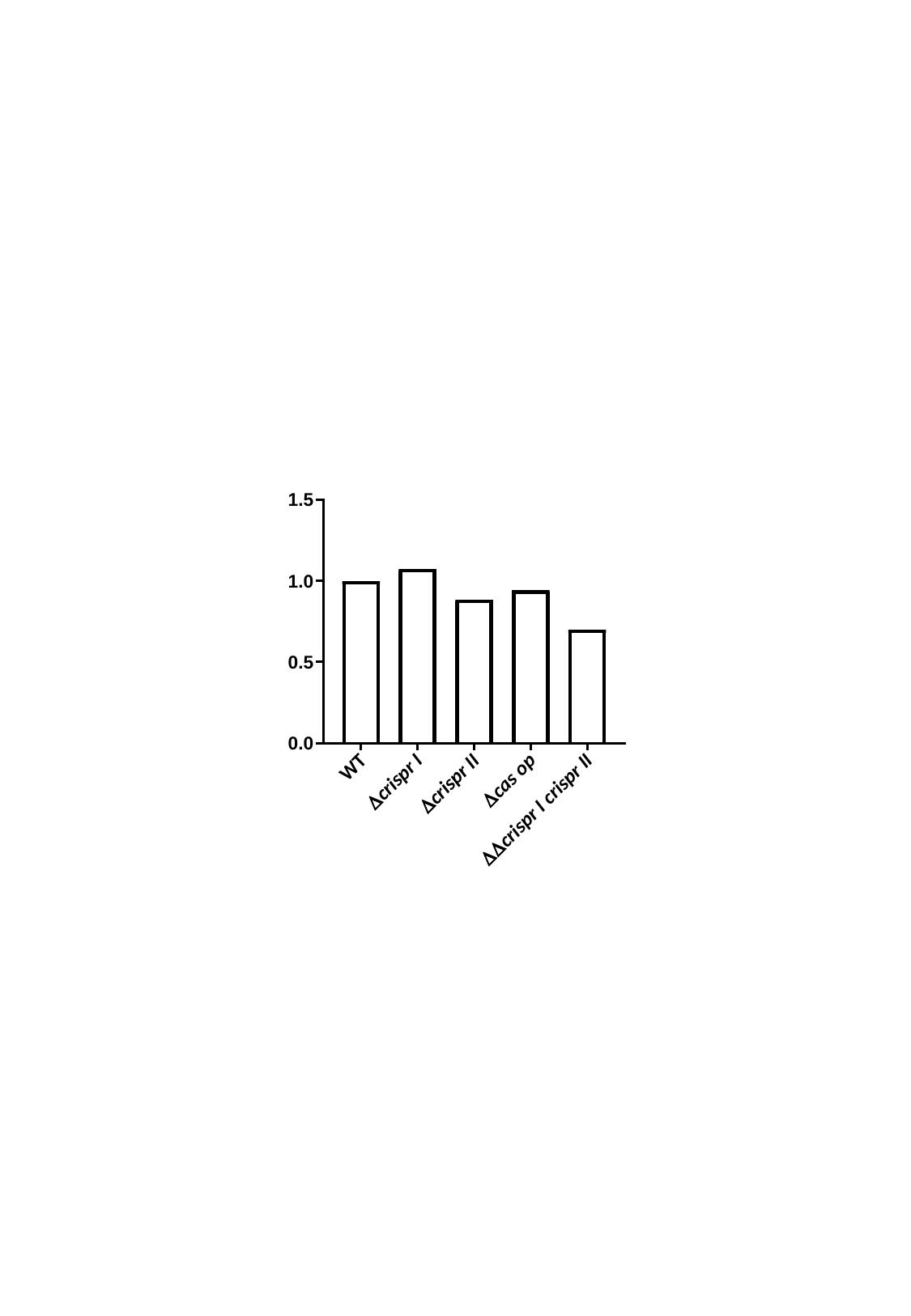

#### Slide 4
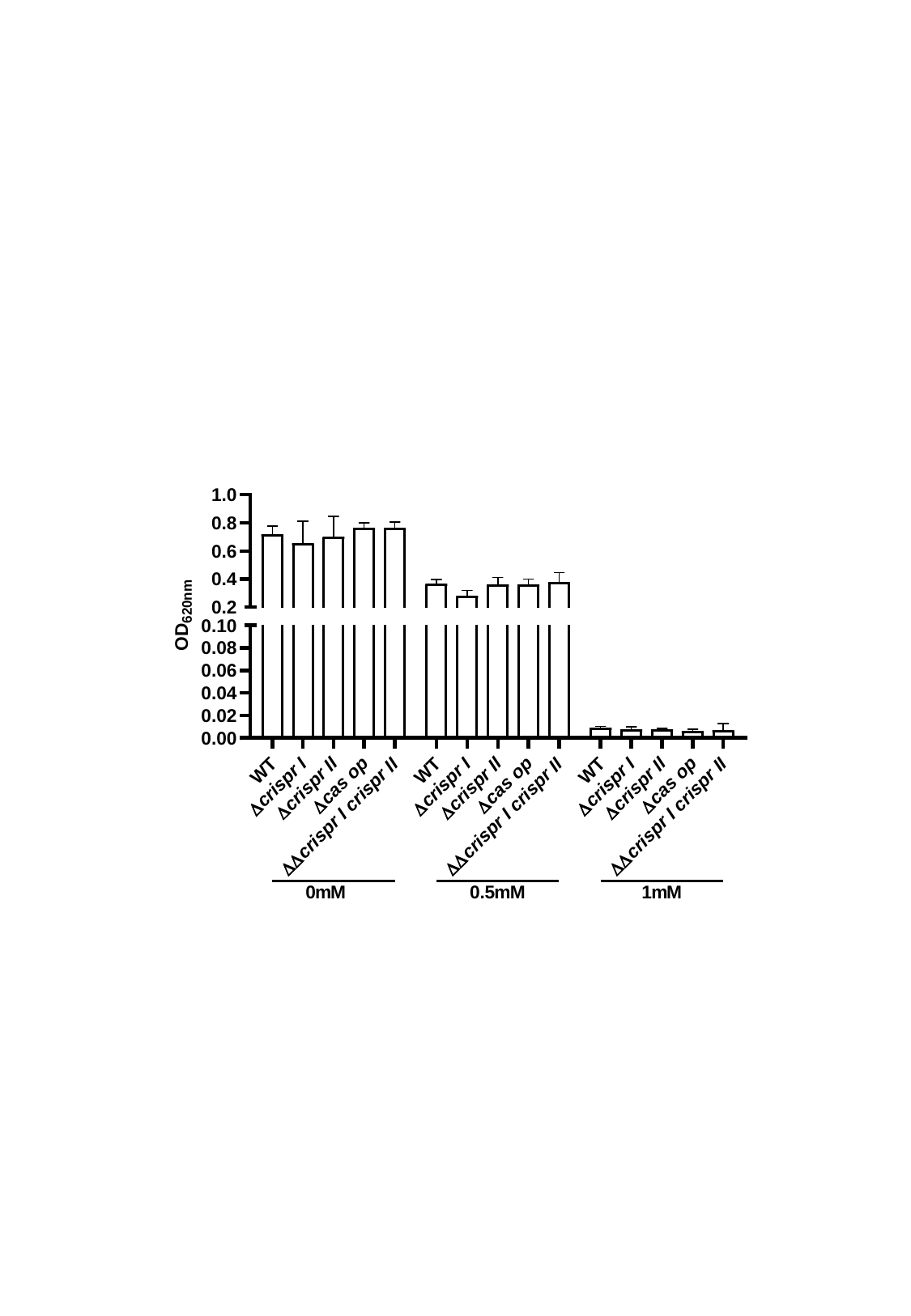

#### Slide 5
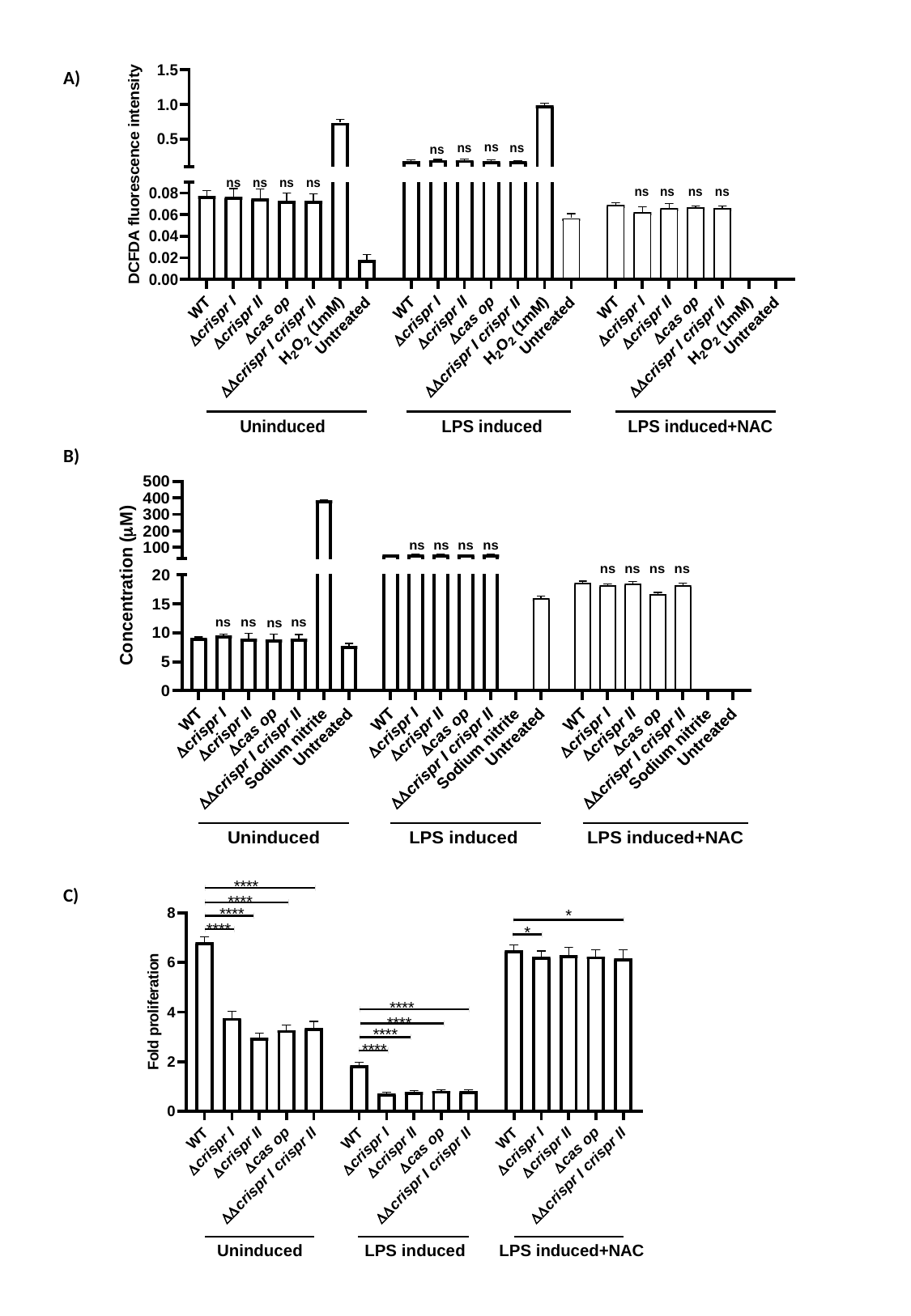

A)
B)
C)

#### Slide 6
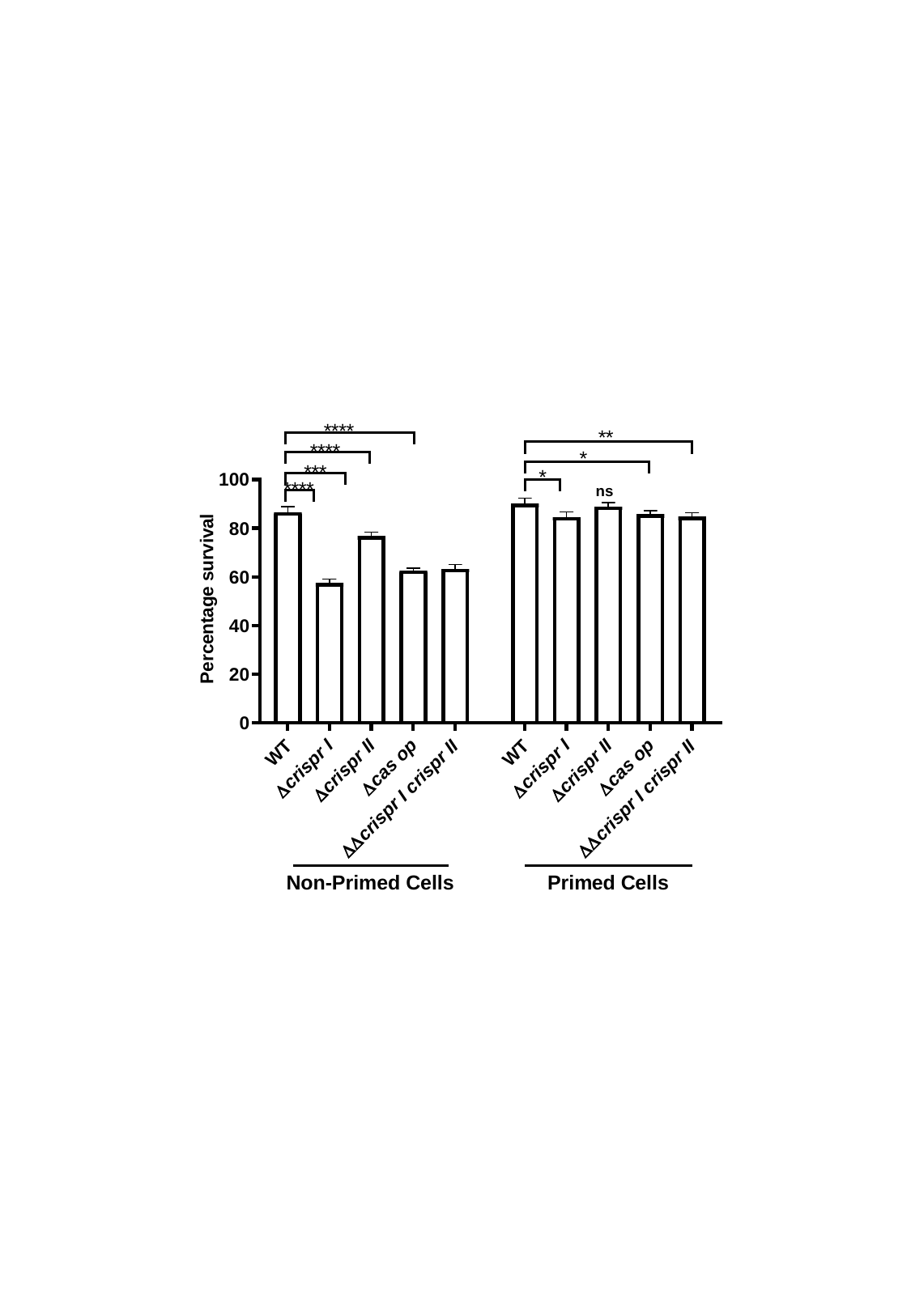

#### Slide 7
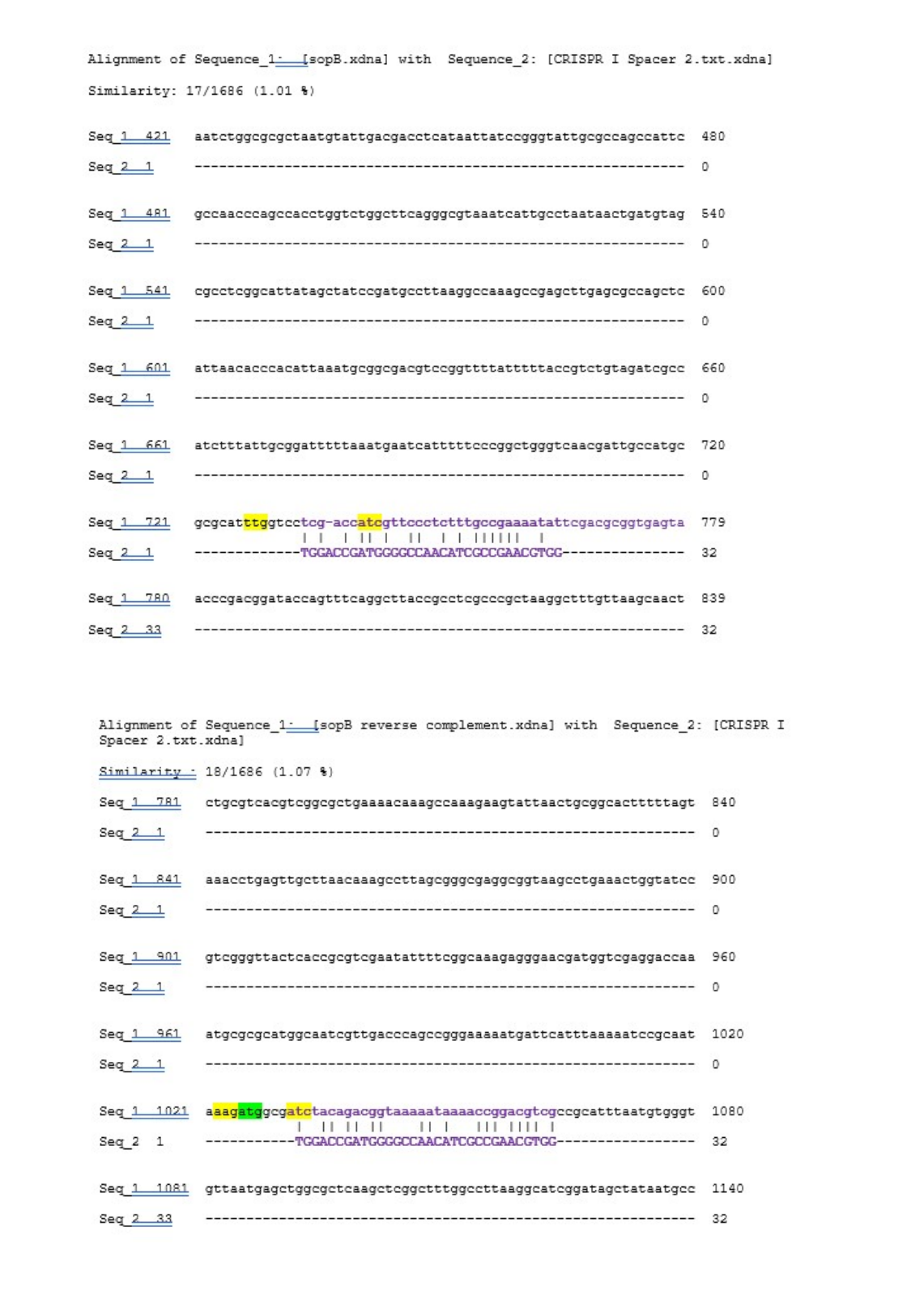

#### Slide 8
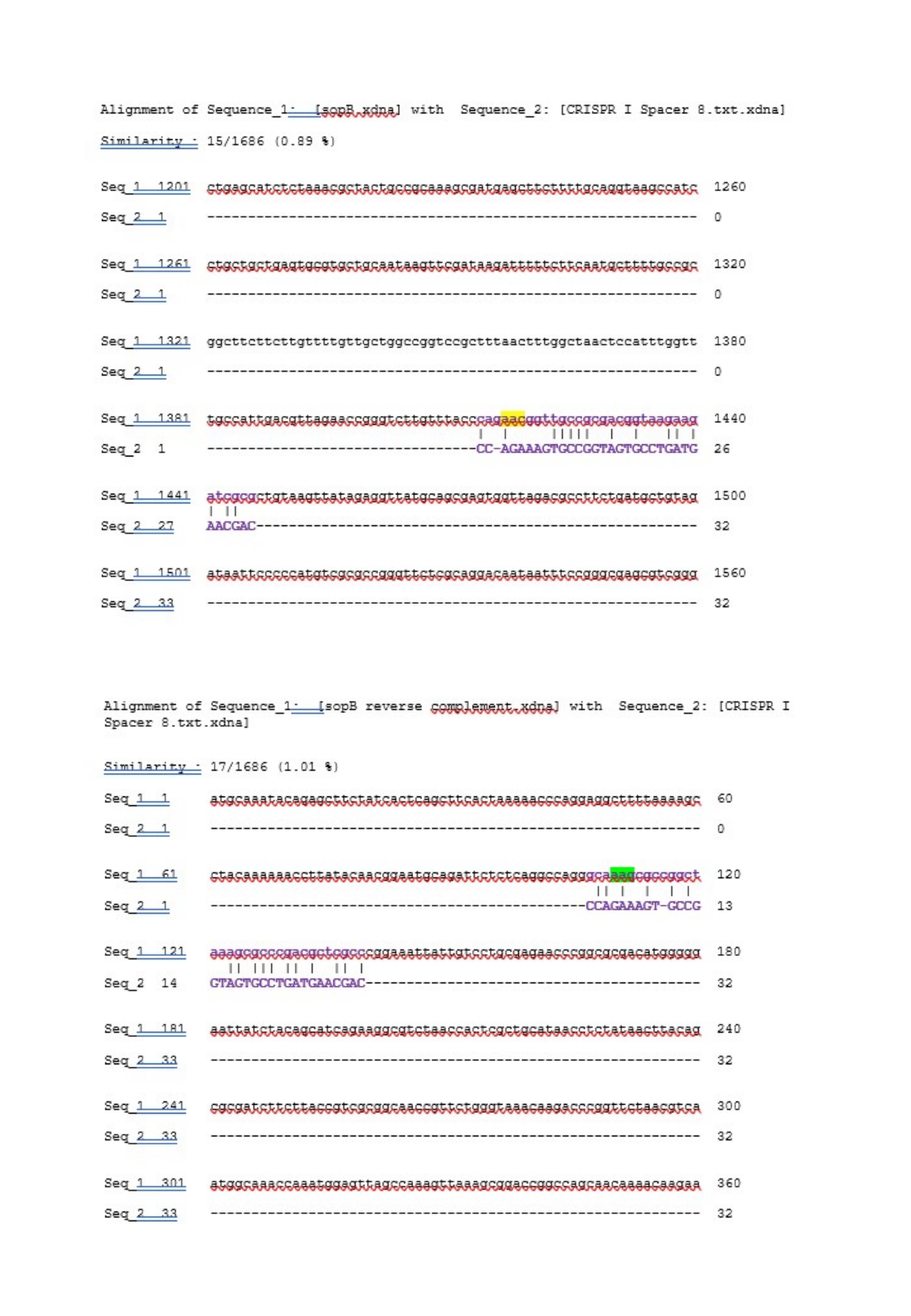

#### Slide 9
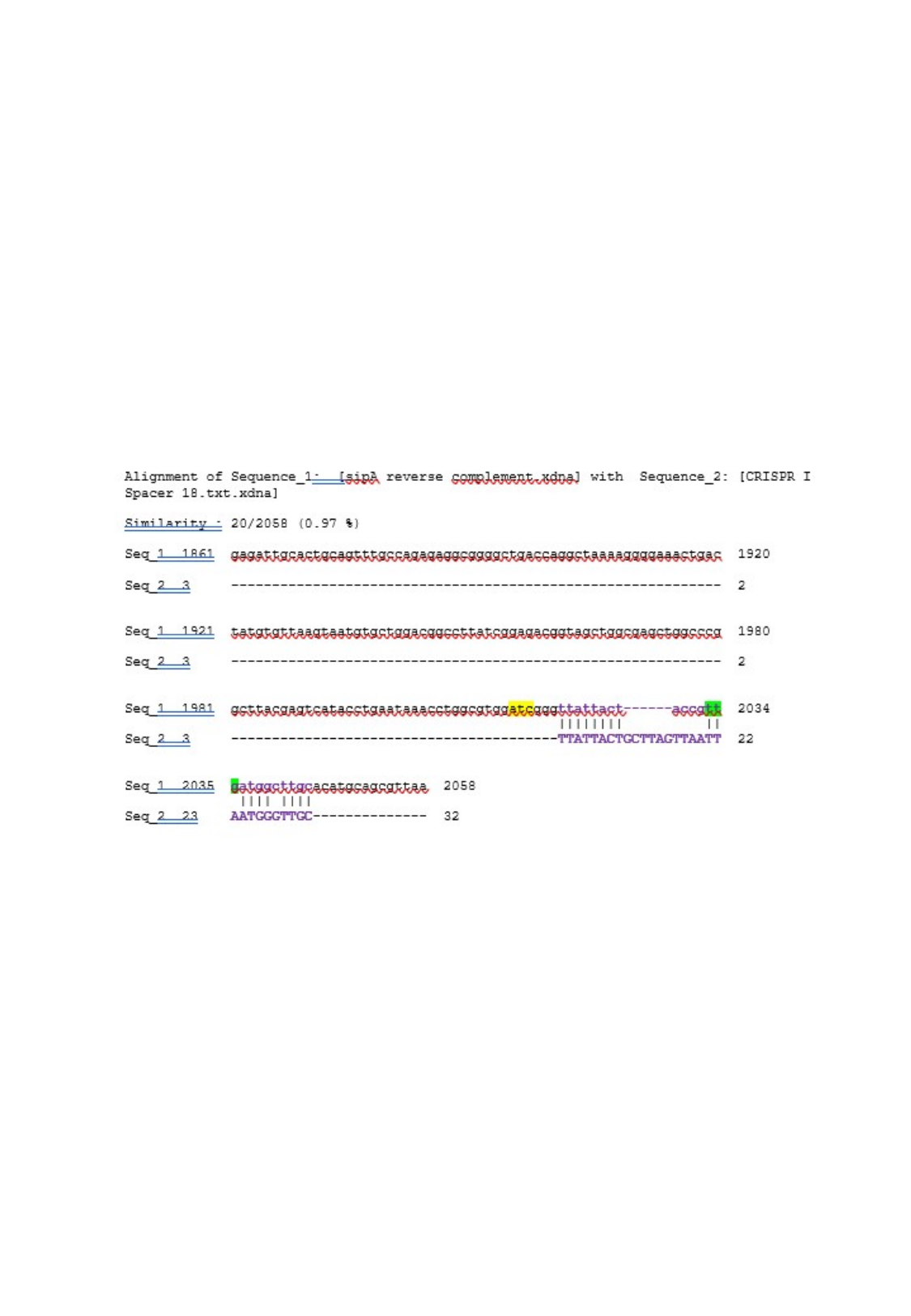

#### Slide 10
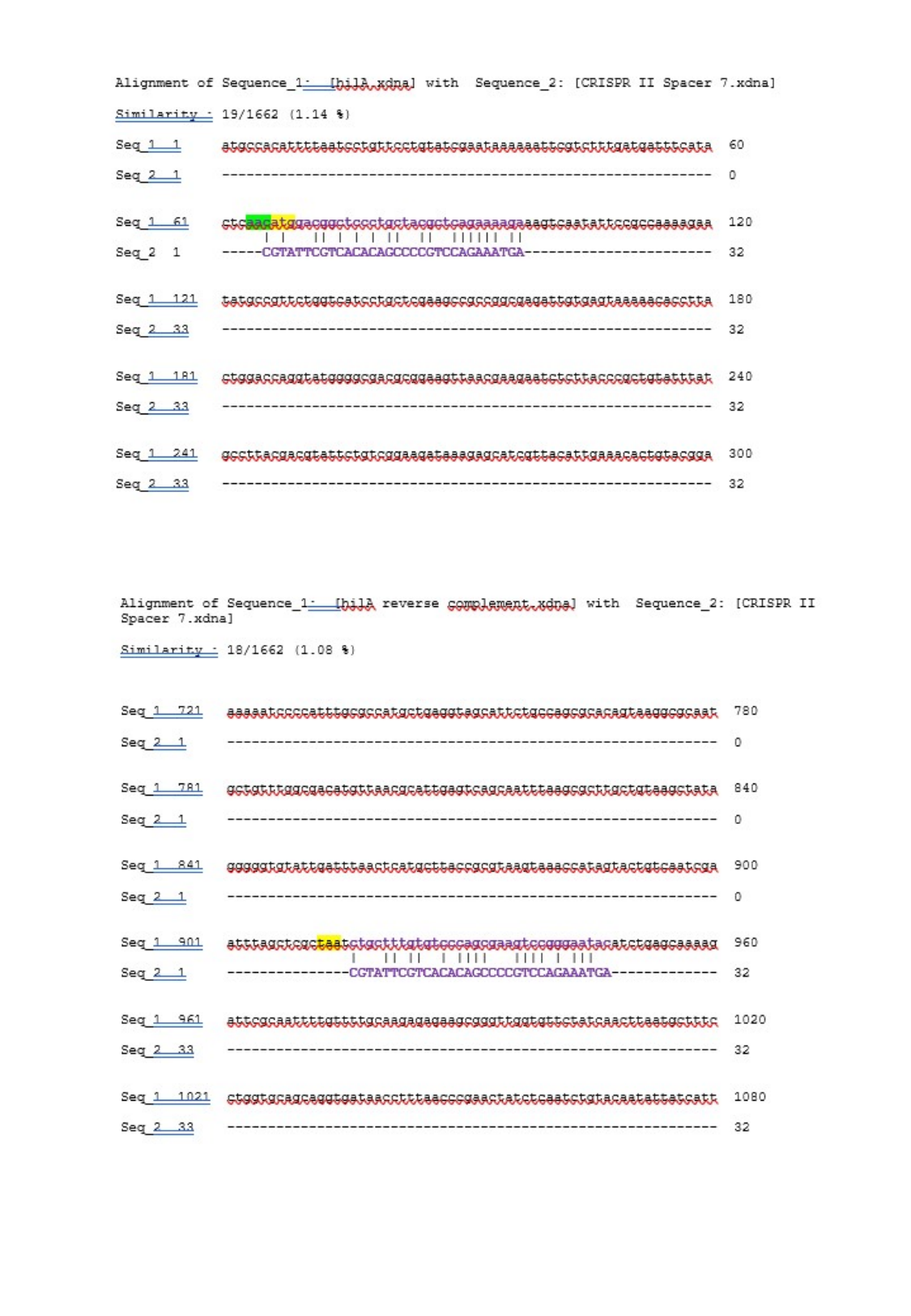

#### Slide 11
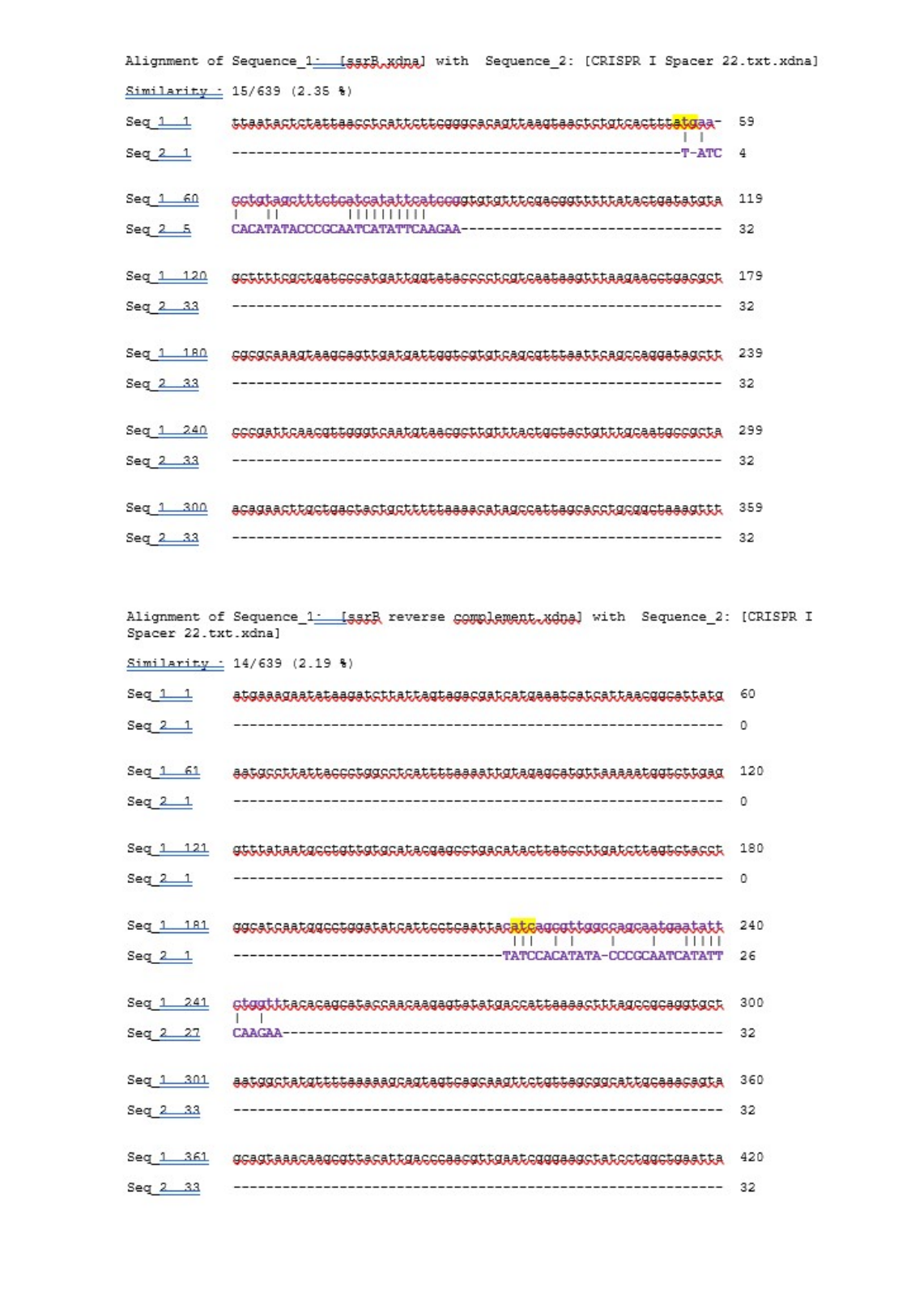

#### Slide 12
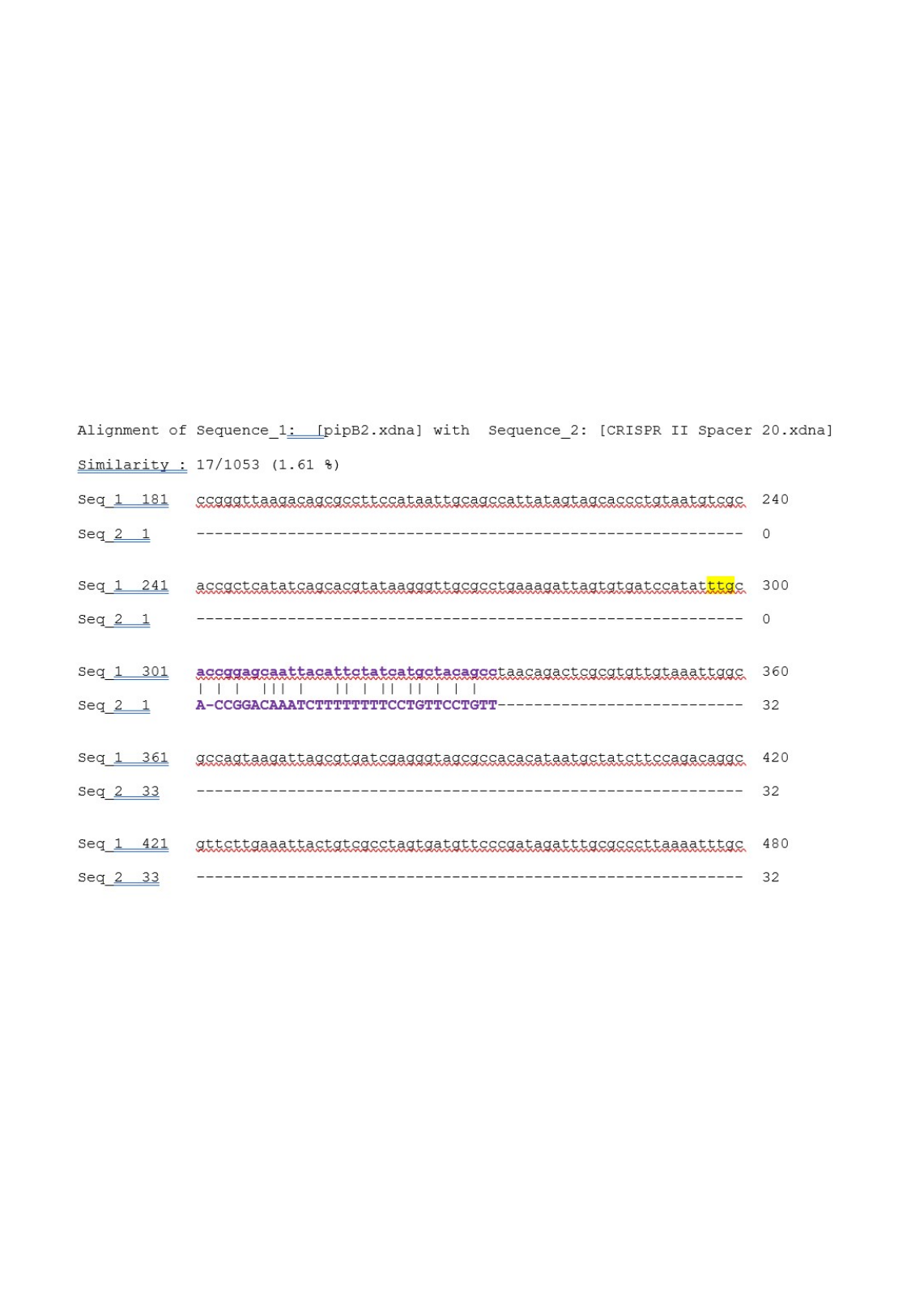

#### Slide 13
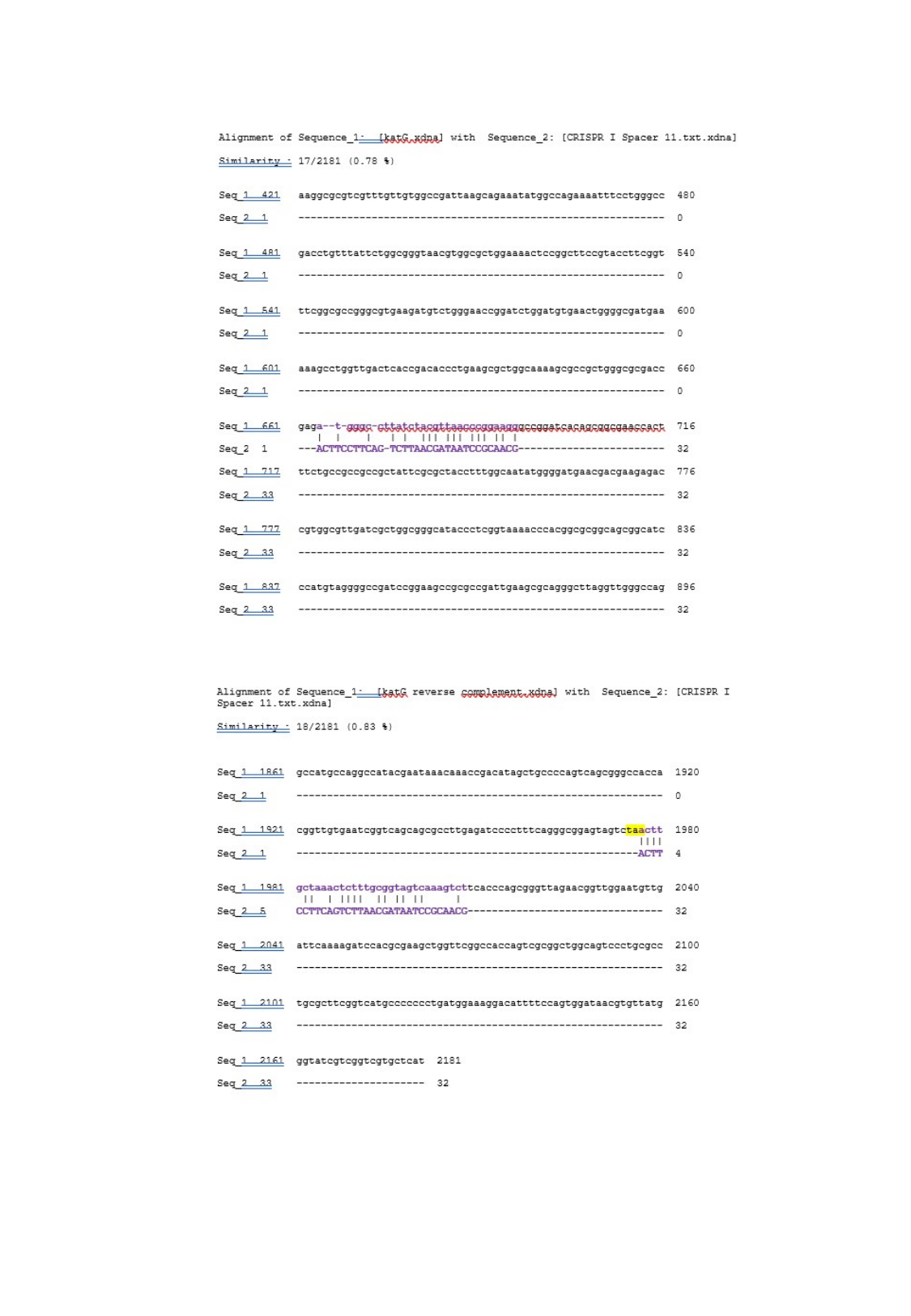

#### Slide 14
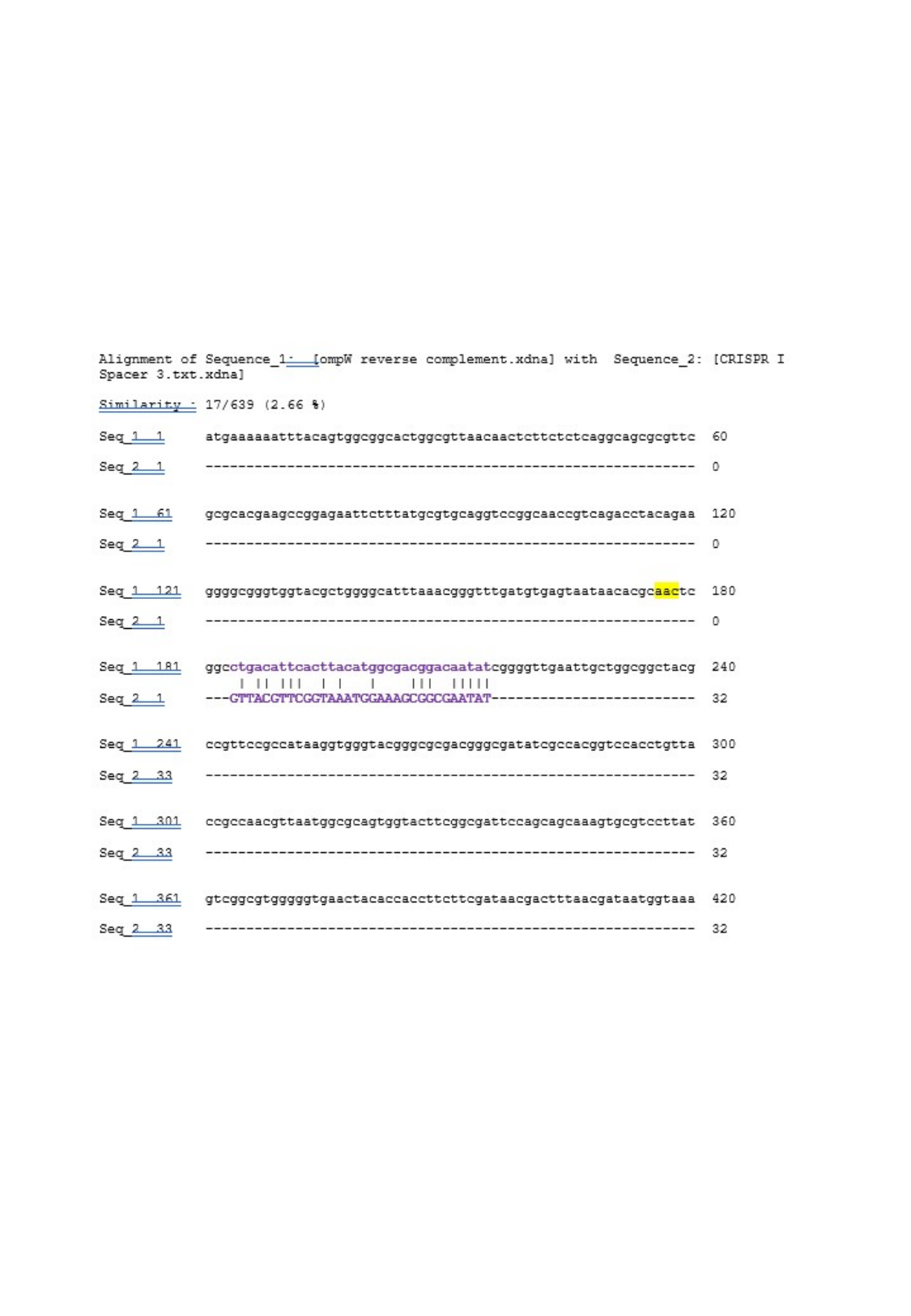

#### Slide 15
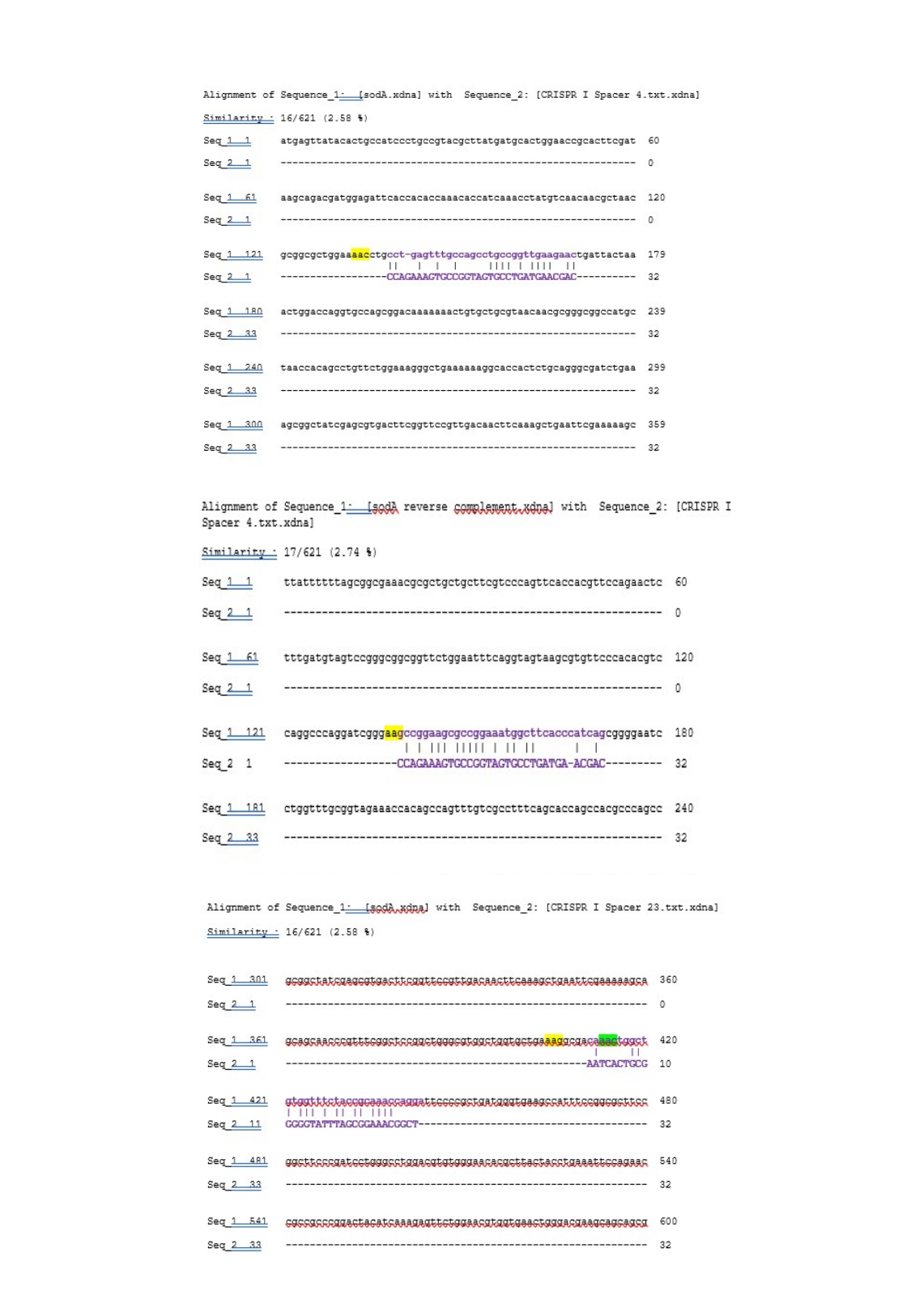
