## Supplementary Table for "Clustered Regularly Interspaced Short Palindromic Repeats-Cas system regulates *Salmonella* virulence"

**Supplementary Table1: Bacterial strains used in the study**

| **Bacterial Strain** | **Genotype and Characteristics** | **Source/ref** |
| --- | --- | --- |
| *Salmonella enterica* serovars Typhimurium 14028s | WT 14028s | A kind gift from Prof. Dipshikha Chakravortty, Indian Institute of Science, India |
| Δ*crisprI* | WT 14028s Δ*crisprI*:: Chl (Chl^r^) | This study |
| Δ*crisprII* | WT 14028s Δ*crisprII*:: Chl (Chl^r^) | This study |
| Δ*cas op* | WT 14028s Δ*cas operon*:: Chl (Chl^r^) | This study |
| ΔΔ*crisprI crisprII* | WT 14028s Δ*crisprI*:: Kan :Δ*crisprI*::Chl (Kan^r^, Chl^r^) | This study |
| *ΔsipD* | WT 14028s Δ*sipD*::Kan (Kan^r^) | A kind gift from Prof. Dipshikha Chakravortty, Indian Institute of Science, India |
| *E.Coli-*OP50 |  | A kind gift from Vidya Devi Negi, Indian Institute of Science Education and Research, Mohali |
| pFPV*-*mCherry | Amp^R^ | A kind gift from Prof. Dipshikha Chakravortty, Indian Institute of Science, India |
| WT-mCherry | WT 14028s transformed with pFPV*-*mCherry Vector (Amp^R^) | This study |
| Δ*crisprI -*mCherry | Δ*crisprI* transformed with pFPV*-*mCherry Vector (Amp^R^) | This study |
| Δ*crisprII-*mCherry | Δ*crisprII* transformed with pFPV*-*mCherry Vector (Amp^R^) | This study |
| Δ*cas op-*mCherry | Δ*cas op* transformed with pFPV*-*mCherry Vector (Amp^R^) | This study |
| ΔΔ*crisprI crisprII-*mCherry | ΔΔ*crisprI crisprII* transformed with pFPV*-*mCherry Vector (Amp^R^) | This study |
| *E.Coli* OP50-mCherry | *E.Coli* OP50  transformed with pFPV*-*mCherry Vector (Amp^R^) | This study |
| Δ*crisprI* +p*crisprI* | Δ*crisprI* complemented with functional CRISPR I array cloned in pQE60(Amp^R^) | This study |
| **Bacterial Strain** | **Genotype and Characteristics** | **Source/ref** |
| Δ*crisprII* +p*crisprII* | Δ*crisprII* complemented with functional CRISPR II array cloned in pQE60(Amp^R^) | This study |

**Supplementary Table2: Sequences of Primers used in the study**

| **Sl. No.** | **Primer Name** | **Nucleotide Sequence** |
| --- | --- | --- |
| 1 | *rpoD* (Forward) | 5’ACATGGGTATTCAGGTAATGGAAGA3’ |
| 2 | *rpoD* (Reverse) | 5’CGGTGCTGGTGGTATTTTA3’ |
| 3 | *hilD* (Forward) | 5’GACGAACCTGGGATGTTG3’ |
| 4 | *hilD* (Reverse) | 5’CGAAATCCATGTGGCCATTG3’ |
| 5 | *hilA* (Forward) | 5’TAACATGTCGCCAAACAGC3’ |
| 6 | *hilA* (Reverse) | 5’GCAAACTCCCGACGATGTAT3’ |
| 7 | *sipA* (Forward) | 5’ATGCGGGAAAGACGCTGA3’ |
| 8 | *sipA* (Reverse) | 5’TCGCCTCAGGAGAATCACTG3’ |
| 9 | *sipD* (Forward) | 5’ATTATCGCAGGCGGCTACTAA3’ |
| 10 | *sipD* (Reverse) | 5’CATTCAGGCTGCTGGTCAAC3’ |
| 11 | *sopB* (Forward) | 5’CGCTCGCCCGGAAATTATTG3’ |
| 12 | *sopB* (Reverse) | 5’GAGGTTATGCAGCGAGTGGT3’ |
| 13 | *ssrB* (Forward) | 5’CCTTATTACCCTGGCCTCA3’ |
| 14 | *ssrB*(Reverse) | 5’CCATTGATGCCAGGTAGACT3’ |
| 15 | *H-NS* (Forward) | 5’ACATCCGTACTCTTCGTG3’ |
| 16 | *H-NS*(Reverse) | 5’ACGAGTGCGTTCTTCCAC3’ |
| 17 | *pipB2* (Forward) | 5’GGATACGTCGGAGAAATGAA3’ |
| 18 | *pipB2* (Reverse) | 5’ATTCGACACGACACCCA3’ |
| 19 | *spiC* (Forward) | 5’TCAGGGCCGAAGGTAATAGC3’ |
| 20 | *spiC* (Reverse) | 5’GGTGTGCTGCAAGCAGTAGT3’ |
| 21 | *sodA* (Forward) | 5’ACCTGCCTGAGTTTGCC3’ |
| 22 | *sodA* (Reverse) | 5’GTTGTTACGCAGCACAGT3’ |
| 23 | *sodCI* (Forward) | 5’ATCACAGTTTCAGAGACACC3’ |
| 24 | *sodCI* (Reverse) | 5’TTCCCGGCATACAACTTG3’ |
| 25 | *katG* (Forward) | 5’GTGAAGATGTCTGGGAACC3’ |
| 26 | *katG* (Reverse) | 5’CCCTTCCGGGTTAACGTA3’ |
| 27 | *ahpC* (Forward) | 5’TTGCCCGACTGAACTGG3’ |
| 28 | *aphC* (Reverse) | 5’GTGCGTGAAGTGAGTATCG3’ |
| 29 | *mgtC* (Forward) | 5’TCTGAGCTCCATGACGAC3’ |
| 30 | *mgtC* (Reverse) | 5’ATCCCTTCGCGCATGAT3’ |
| 31 | *ompW*(Forward) | 5’CGGGTTTGATGTGAGTAATAAC3’ |
| 32 | *ompW*(Reverse) | 5’GAAGTACCACTGCGCCATT3’ |
| **Sl. No.** | **Primer Name** | **Nucleotide Sequence** |
| 33 | *pmrD*(Forward) | 5’ GCGTGCCATGTTCTGGT3’ |
| 34 | *pmrD*(Reverse) | 5’ CAGGATATCGCCGGGACT3’ |
| 35 | *pmrA*(Forward) | 5’ CGTGTGATGGCGTTTCG3’ |
| 36 | *pmrA*(Reverse) | 5’ TCTGTCGGATTCGCGTCA3’ |
| 37 | *pmrH*(Forward) | 5’ CGATACGCTGATGGTCACG3’ |
| 38 | *pmrH*(Reverse) | 5’ CTTCAATCACTGGAATGCCA3’ |
| 39 | *pmrE*(Forward) | 5’ ATCAACCCCGCAGCAGT3’ |
| 40 | *pmrE*(Reverse) | 5’ ATCACGATACGGGAGGGATA3’ |
| 41 | *pag*D(Forward) | 5’CAGTTCAGGCCATTGTTCT3’ |
| 42 | *pagD*(Reverse) | 5’ATCAGTATTTGTAGGTGGTTGC3’ |
| 43 | *pagB*(Forward) | 5’TGATTGCCGTGAGCGT3’ |
| 44 | *pagB*(Reverse) | 5’CACCATTACGCCTGCGA3’ |
